# Quantification of Cas9 binding and cleavage across diverse guide sequences maps landscapes of target engagement

**DOI:** 10.1101/2020.09.09.290668

**Authors:** Evan A Boyle, Winston R Becker, Hua B Bai, Janice S Chen, Jennifer A Doudna, William J Greenleaf

**Affiliations:** Department of Genetics, Stanford University School of Medicine, Stanford, CA 94305, USA; Program in Biophysics, Stanford University, Stanford, CA 94305, USA; Department of Molecular and Cell Biology, California Institute for Quantitative Biosciences (QB3), University of California, Howard Hughes Medical Institute, Department of Chemistry, and the Innovative Genomics Institute, University of California, Berkeley, Berkeley, United States; MBIB Division, Lawrence Berkeley National Laboratory, Berkeley, United States; Gladstone Institutes, University of California, San Francisco, San Francisco, United States; Department of Applied Physics, Stanford University, Stanford; Department of Cellular & Molecular Medicine, University of California San Diego, San Diego, CA; Biological Sciences graduate program, University of California San Diego, San Diego, CA; Mammoth Biosciences, Inc., South San Francisco, CA

## Abstract

The RNA-guided nuclease Cas9 has unlocked powerful methods for perturbing both the genome through targeted DNA cleavage and the regulome through targeted DNA binding, but limited biochemical data has hampered efforts to quantitatively model sequence perturbation of target binding and cleavage across diverse guide sequences. We present scalable, sequencing-based platforms for high-throughput filter binding and cleavage, then perform 62,444 quantitative binding and cleavage assays on 35,047 on- and off-target DNA sequences across 90 Cas9 ribonucleoproteins (RNPs) loaded with distinct guide RNAs. We observe that binding and cleavage efficacy, as well as specificity, vary substantially across RNPs; canonically studied guides often have atypically high specificity; sequence context surrounding the target significantly influences Cas9 on-rate; and Cas9 RNPs may sequester targets in nonproductive states that contribute to “proofreading” capability. Finally, we distill our findings into an interpretable biophysical model that predicts changes in binding and cleavage for diverse target sequence perturbations.

## Introduction

*Streptococcus pyogenes* (Spy) Cas9 has been widely adopted as a platform for perturbing gene expression and protein levels in human cells (*1*). In this type II CRISPR system, the CRISPR associated protein Cas9 performs targeted search and cleavage of double-stranded DNA guided by a CRISPR RNA (crRNA) that is complementary to the target sequence. This native CRISPR-Cas9 bacterial system has also been engineered to bind to DNA without inducing cleavage, creating a powerful platform for modulating gene expression by fusing activators or repressors to catalytically inactive Cas9 (dCas9) then targeting binding to specific genomic loci(*2*).

Ideal gene editing or modulation tools require both high levels of sensitivity (i.e. high probability of binding or cleavage at a targeted site) as well as excellent specificity (i.e. low probability of binding or cleavage at non-targeted sites) (*3*, *4*). Because the biophysical processes involved in target search and binding necessarily underlie this sensitivity and specificity, they have been the subject of extensive investigation. Such work has revealed that the Cas9 RNP first associates to an NGG Protospacer Adjacent Motif (PAM), then hybridizes to 8–12 target nucleotides located next to this PAM, known as the “seed” region. Mismatches within this seed region inhibit stable RNP:target complex formation, whereas mismatches located distal to this region act to reduce the lifetime of RNP:target complexes (*5*). Building off of this work, and in combination with insights gleaned from characterizing Cas9 structures (*6*–*8*), others have characterized how DNA unwinding and subsequent conformational changes gate the distinct domains responsible for catalytic cleavage (HNH and RuvC) after binding (*9*–*11*). Finally, recent work has suggested that Cas9 RNP:target interactions proceed along multiple paths, some of which appear to pass through or end in nonproductive states that act as sinks (*10*, *12*, *13*).

Thus, while the steps of canonical Cas9 binding are known, principles underlying sequence-dependent efficacy across guide sequences and sequence-dependent sensitivity to sgRNA:target mispairing given a guide sequence have been less comprehensively addressed. Most biophysical studies have measured relatively few RNP:target pairs, and while recent work has extended the number of off-target binding measurements per guide, the total number of sgRNAs profiled remains limited (*14*, *15*). Further, even scalable technologies for measuring DNA-protein interactions, such as HiTS-FLIP (*16*), HT-SELEX (*17*), Bind-n-seq (*18*), and BunDLE-seq (*19*), often have limited kinetic resolution and are ill-suited to measuring either transient or low affinity interactions, making characterization of weak off-target interactions problematic. The lack of diverse biophysical data across many guides and many off-target sites leaves few avenues for modeling Cas9 off-target activity (*20*, *21*).

To measure Cas9 binding in a quantitative and scalable manner, we developed a massively parallel nitrocellulose filter binding assay by replacing autoradiography with a sequencing based readout, enabling a label-free measurement of dCas9 RNP binding kinetics to thousands of off-targets in a single experiment (*15*). Here we further optimize and parallelize this filter-binding technique, and generate binding and cleavage data on over 45,000 on- and off-target DNA sequences across 91 distinct sgRNAs. In so doing, we more than double the number of publicly available off-target binding measurements. Our data highlight the diversity of RNP biochemical behavior when loaded with different sgRNAs: some sgRNAs are highly specific and exhibit large changes in binding when mismatches are present at the concentrations probed, while others are much less sensitive to mismatches. We demonstrate that context sequence outside the target and PAM can significantly modulate RNP association rates, which are correlated with Cas9 targeting efficacy in cells. Finally, we develop a predictive biophysical model for Cas9 binding and cleavage of off target sites.

## Results

### Massively parallel filter binding enables scalable, quantitative measurement of Cas9 binding

We first selected 90 guide RNA sequences and designed a matched library of approximately 600 DNA targets for each of these guides (one “sublibrary”). Each sublibrary included DNA targets with all single mismatches, 66 contiguous double mismatches, 10 noncontiguous double mismatches, all single RNA:DNA bulges plus select double and triple bulges, 230 contiguous mismatch series consisting of rA:dA, rC:dC, rG:dG and rU:dT mismatches from a start to end position, and control sequences common to all sublibraries. In total, 54,349 distinct targets were designed (Supplementary Table 1). For each sublibrary, a corresponding sgRNA was prepared and loaded in dCas9 while the DNA was split and PCR barcoded with 16 distinct timepoint primers (Figure 1A) that allowed quantification of a binding time course (see methods).

**Figure 1:**
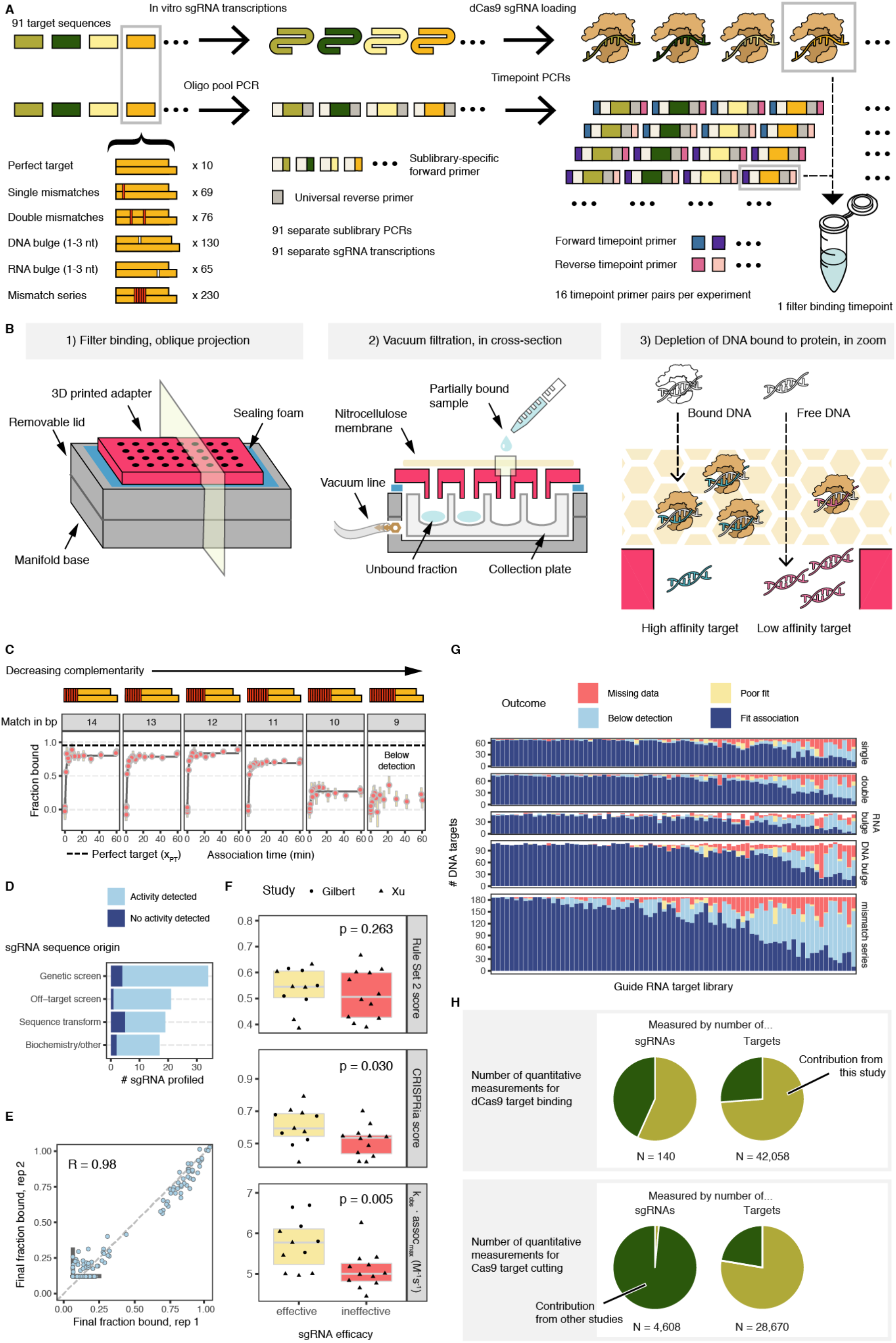
Kinetic profiling of dCas9 binding by massively parallel filter binding. A) Experimental overview. Cas9 targets designed to match 90 sgRNAs are synthesized on an oligonucleotide array. Using 91 distinct sublibrary primers, each sgRNA’s targets are amplified separately. Each library is barcoded in a second PCR with 16 pairs of forward and reverse indices. Following filter binding, all timepoints and sublibraries in the experiment are pooled and sequenced in one run. B) Three views of filter binding apparatus mid-experiment: 1) an adaptor rests on top of a 96-well plate sized vacuum manifold; 2) sample is passed through a nitrocellulose membrane placed on top of the adaptor and collected in a deep 96-well plate below; 3) as sample passes through the nitrocellulose, bound DNA is trapped in the membrane while free DNA passes through unimpeded. C) Example association curves fit from count data per target sequence. Each panel shows varying complementarity to λ1 sgRNA starting with matching nucleotides from the PAM-proximal end. Yellow bars signify 90% confidence intervals. 9 bp of complementarity and below showed low final binding levels and were classified as ‘below detection.’ The dashed line signifies the final fraction bound of the perfect target (20 bp matching). D) Summary of sgRNAs included in study including how the sgRNAs were curated and whether they showed evidence of binding. E) Reproducibility of λ1 targets’ final fraction bound across two sublibrary amplifications and sgRNA preparations. Bold gray lines in lower left denote limits of detection. F) sgRNAs nominated as effective or ineffective by genetic screening data exhibit greater difference in their empirical perfect target productive on-rates than scores from computational models for sgRNA activity such as ‘Rule Set 2’ and ‘CRISPRia.’ G) Summary of association curves for all studied sgRNA sublibraries. Low counts and sequence bias cause missing data. Sequences showing little or no depletion over the course of the experiment are deemed ‘below detection.’ Fits that produce estimates beyond the dynamic range of the experiment are binned as ‘poor fits.’ H) Pie chart demonstrating relative scale of presented data compared to other published studies of Cas9 off-target activity.

We next designed a massively parallel filter-binding apparatus to permit processing binding timecourses of sublibraries in 96-well plates (Figure 1B). As part of this workflow, we use a nitrocellulose membrane to bind to protein-bound DNA targets, then collect and sequence the unbound DNA in the flow-through, thereby quantifying binding through measurement of depletion. The nitrocellulose membrane is placed over a 3D-printed 96-pronged adapter engineered to mate with a 96-deep well plate. When the specified association time for a sublibrary timepoint elapses, the sample is applied to the membrane and flow-through collected by vacuum filtration. At the end of the experiment, a fraction of this flow-through is taken from each well to be pooled and sequenced. Relative to our previous protocol (*15*), this 96-well design required 70% less hands-on time, 90% less reaction volume, and 85% less expense.

Raw sequencing data was fit to a one-step association model (see methods), resulting in two fit parameters: final fraction bound (f_final_) and an observed rate (k_obs_). Confidence intervals for each timepoint were constructed assuming Poisson noise. Targets for which the first timepoint neared the binding level of the last timepoints were fit to final fraction bound only. Furthermore, targets wherein timepoints’ confidence intervals for estimated fraction bound overlapped zero were separately flagged as binding below our detection limit (see methods). Off-targets with an extreme fit rate or final fraction bound were flagged as poor fits. Some targets, particularly DNA targets with low representation in the library, could not be fit and represent missing data.

We first conducted filter binding association experiments on all 91 sublibraries with 5 nM dCas9 RNP and 100 pM total DNA. Overall, fit lines overlapped count-based confidence intervals and agreed well with past work on dCas9 sequence specificity, such as the finding that a 5-10 bp seed sequence is sufficient for binding at 5 nM RNP (*22*, *23*) (Figure 1C). 12 of 91 (13%) perfect target sequences could not be fit, largely because binding levels were below the threshold of detection. Observation of detectable binding activity depended on the origin of the curated sequence (Figure 1D). Half of the unfit sgRNAs contained 17 or more guanines/cytosine base pairs compared to 13% of fit sgRNAs (p = 2e-3, binomial test). By screening sgRNA sequences in silico with RNAfold (*24*), another 4 exhibited extensive secondary structure (Figure S1A) that could interfere with the folding of sgRNA hairpins, a characteristic known to lead to poor sgRNA performance (*25*). Across the 79 sgRNAs with valid perfect target measurements, binding to 29,232 target sequences was quantified (Figure 1G; Supplementary Table 2), substantially more off-target measurements than previous efforts (Figure S1B and Figure 1H). An additional 5,983 targets were classified as binding below our detection limit. Amongst these, the targets least likely to be quantified were predicted to bind with RNA bulges or with a long series of mismatched bases. To quantify experimental variability, we prepared and assayed two separate sublibraries for the λ phage genome target known as λ1. We found that the fit final fraction bound was in good agreement across replicates (R = 0.98, Figure 1E).

We next aimed to compare the parameters estimated from these experiments with *in vivo* activity scores. We first classified the subset of sgRNAs with published CRISPRi activities as either effective or ineffective (*20*, *21*) and assessed whether the two classes exhibited differences in fit biophysical parameters. As a baseline, we applied two published predictive algorithms for CRISPRi guide activity: Rule Set 2 (*3*) and the CRISPRia model scoring (*21*) (Supplementary Table 3). Scores reported by both methods were higher for effective than ineffective sgRNAs (Figure 1F). However, differences in Rule Set 2 scores were not significant, while CRISPRia scores met statistical significance (p = 0.030, Wilcoxon Rank Sums test). We then compared the discriminative power of our quantified k_obs_ and f_final_. Final fraction bound for on-target sequences mostly exceeded 50% and did not correlate with guide efficacy. In contrast, association rates for effective sgRNAs were significantly faster than those of ineffective sgRNAs (p = 0.005, Wilcoxon Rank Sums test). This observation is consistent with recent CRISPRi data demonstrating that apparent association rates govern CRISPRi activity in human cells (*26*).

### High-throughput kinetic measurements reveal diverse sequence landscape of dCas9 association

To assess the sequence specificities of association, we first visualized the distribution of f_final_ and k_obs_ for off-target sequences possessing stretches of zero to twenty complementary nucleotides at the PAM-proximal end of the target (Figure 2A). Observed association rates spanned a 30-fold range across perfect targets, but for a given sgRNA, off-target association rates usually fell within in a narrow range. Most sgRNAs showed little or no decrease in the final fraction bound (at 5 nM loaded Cas9) until complementarity dropped below 12 bp. However, some sgRNAs exhibited large decreases in final fraction bound when as few as 1 or 2 distal mismatches were introduced.

**Figure 2:**
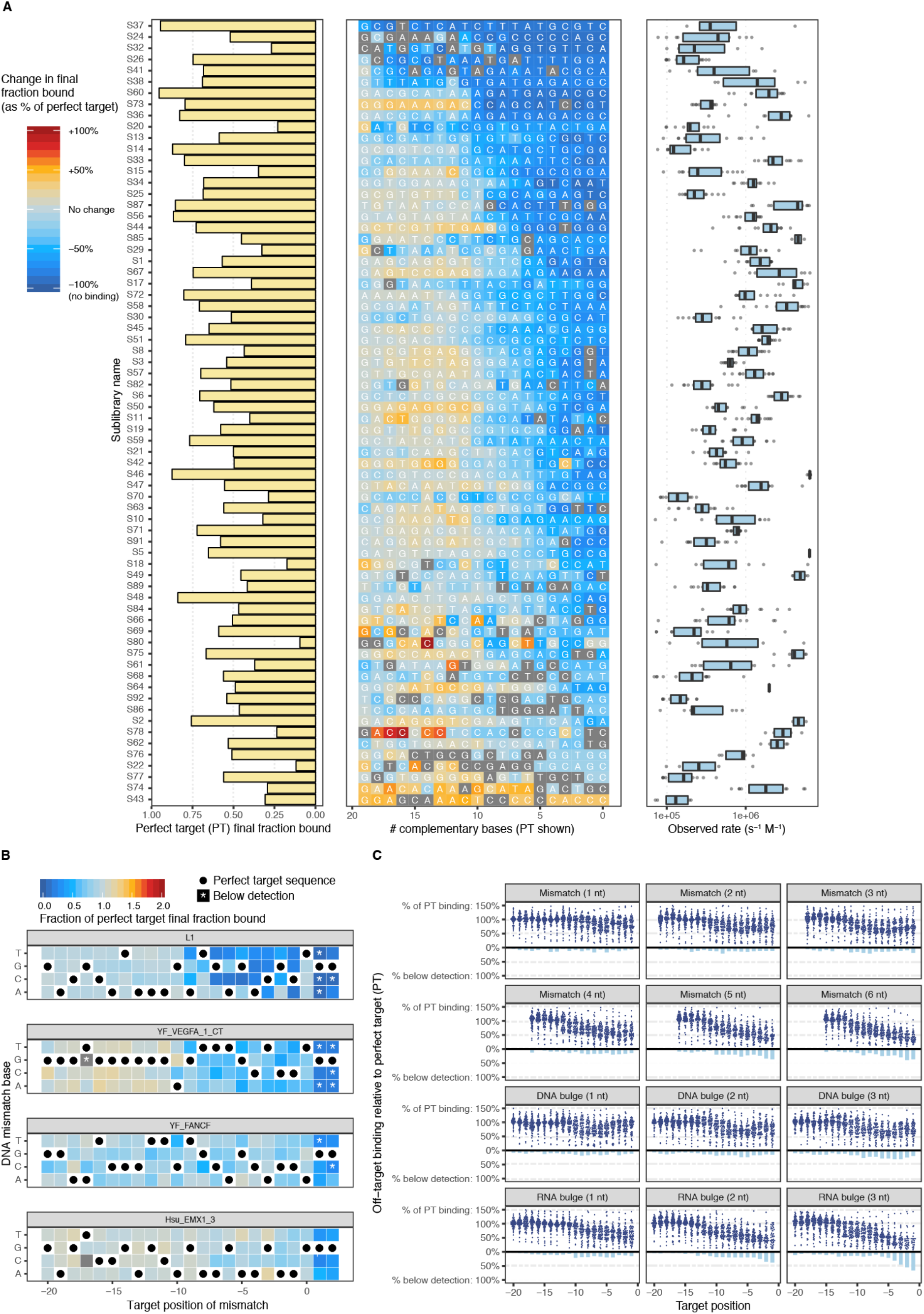
Diversity of dCas9 off target association across sgRNA. A) Association data for sgRNA from zero (PAM-proximal) to perfect (PAM-distal) complementarity. The perfect target final fraction bound is shown for each sgRNA on left; off-target binding relative to perfect target is shown in the middle; the distribution of observed on-rates for the 20 targets in the center column is shown on right. B) Single mismatch data for 4 sgRNAs. PAM mutations are often near or below the limit of detection (asterisks), but many seed mismatches (positions −8 to −1) are within dynamic range. C) Summary of association data collected across measured sgRNAs. Each panel visualizes mismatch or bulge series (of varying size) positioned alongside each base pair of the target sequence. Above the line, binding is reported as a percentage of perfect target binding level. Below the line, bars summarize the percent of sgRNA for which the off-target binding was below detection. For example, the majority of sgRNAs exhibit undetected binding for 3-bp RNA bulges in the seed (bottom right facet).

Although the λ1 sgRNA has been the sgRNA of choice for characterizing Spy Cas9 RNPs, the biophysical properties of the λ1 RNP appear to be atypical: f_final_ and k_obs_ for the λ1 perfect target are among the largest of the sgRNAs we profiled. Additionally, f_final_ for λ1 RNP declines especially steeply for off-targets with fewer than 11 base pairs of complementarity (Figure 2A: sublibrary S60). Most RNPs exhibit final fraction bound that is near perfect target levels until complementarity drops to 8 or 9 base pairs. Declines in final fraction bound for λ1 single mismatch targets are also more extreme than for most sgRNAs. Some, such as FANCF and EMX1 site 3, are minimally perturbed by single mismatches in their targets, unless the mismatches disrupt the canonical PAM (Figure 2B).

Across all sgRNAs, most RNA:DNA mismatches or bulges had small effects on final fraction bound (Figure 2C; Supplementary Table 4). Single RNA:DNA mismatches had particularly modest impact, generally only visible in first 7 bp of the seed. Contrary to expectation, insertion of additional base pairs into the DNA target, which is expected to lead to the formation of a DNA bulge in the RNA:DNA duplex within the RNP, were tolerated nearly as well as mismatches. Further investigation showed that DNA bulges that matched the identity of the PAM-proximal nucleotide fared better than bulges of nucleotides that did not match but were located at the same position (p = 6e-5, one-sided Wilcoxon Rank Sums test). Interestingly, this DNA insertion preference was most prominent at end positions (−1, −18, and −19) or center positions (−8 through −11) (Figure S2A; Supplementary Table 5). Positions outside of these regions did not exhibit such bias whether tested individually or in aggregate. Deletion of DNA target bases, which is expected to lead to the formation of an RNA bulge, typically led to a greater decrease in the final amount of target binding. For 3 bp RNA bulges in the 5 bp nearest the PAM, the majority of off-targets were at or below the limit of detection.

Having characterized off-target binding to naked DNA, we next asked if the final fraction bound (f_final_) for a given sgRNA might be an accurate proxy for CRISPRi silencing capability. We first assessed predicted sgRNA CRISPRi activity for mismatched targets using a recently published model for CRISPRi efficacy at mismatched sites (*26*) by comparing f_final_ measurements to predicted activities for the λ1 sublibrary. We found that the vast majority of off-targets fell into one of two categories: low (<10%) predicted activity and low f_final_ (<30%), or moderate to high (>10%) predicted activity and high f_final_ (> 60%) (Figure S2B) meaning that our biophysical measurements generally agreed with prior modeling efforts (Spearman R = 0.711, p = 4e-23); furthermore, across all sublibraries with at least 50 f_final_ measurements, the correlation between these two metrics were overwhelmingly positive (65 positive / 69 tested, mean Spearman correlation of 0.453; Figure S2C; Supplementary Table 6).

### Cleavage assays highlight persistent sub-saturating activity of Cas9 RNPs

To investigate sequence-dependence of cleavage globally, we used the same barcoded 91 sublibraries to collect timepoint-resolved cleavage data at 5 nM active Spy Cas9 (Figure 3A). Instead of passing samples through a nitrocellulose membrane, samples were quenched with EDTA and heat inactivated prior to sequencing, rendering cleaved products unsequenced. Because sub-libraries were barcoded prior to the cleavage assay, libraries were directly sequenced without PCR amplification, and resulting counts were used to determine the observed cleavage rate and final fraction cleaved (Figure 3B).

**Figure 3:**
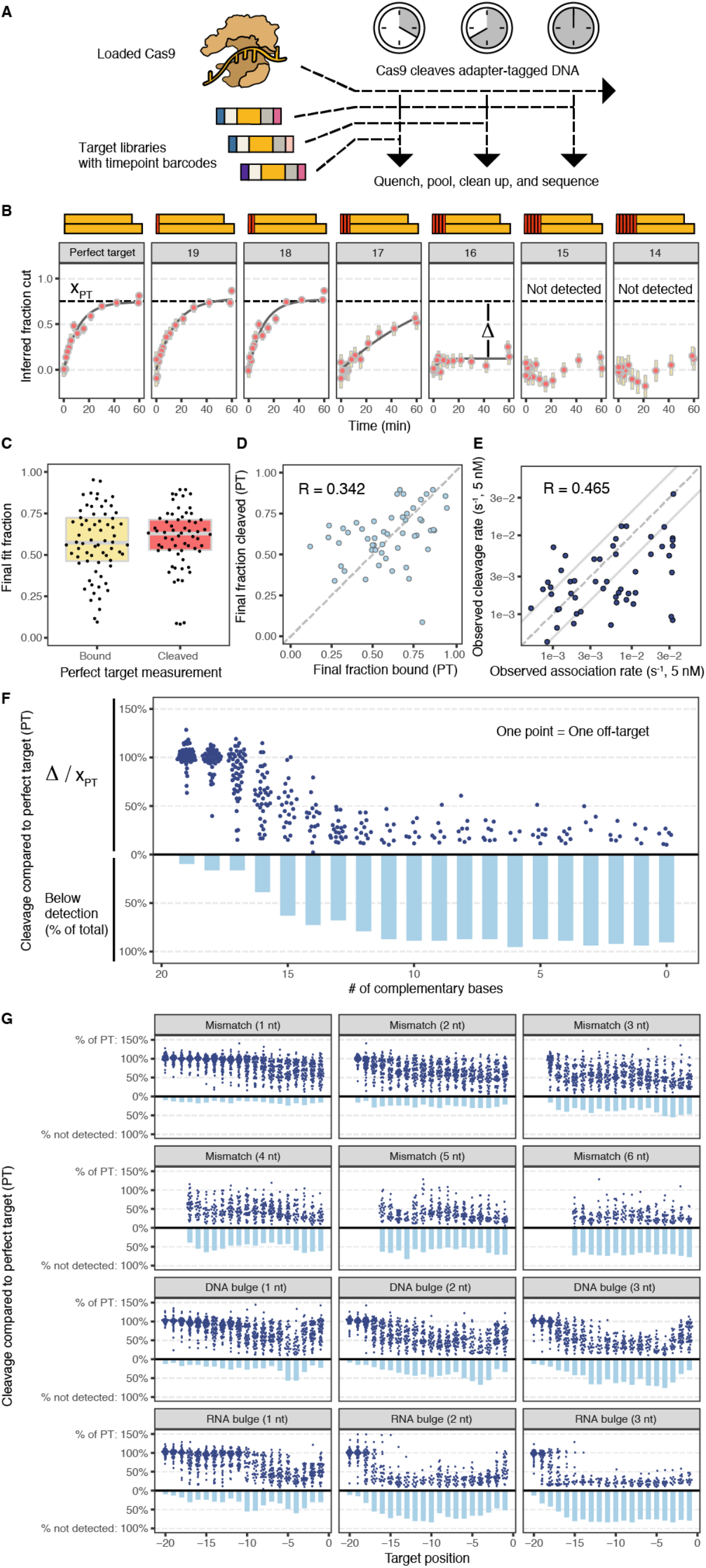
Matched cleavage data for Cas9 off-target libraries. A) Cas9 cleavage experiments consist of timepoints read out in barcoded libraries in a PCR-free sequencing library. B) Example cleavage data demonstrating cleavage as a function of base pairs of complementarity. C) Final binding and cleavage levels for perfect targets are widely distributed. D) Final binding level (x-axis) is moderately correlated with final cleavage level (y-axis) for perfect target sequences. E) Joint distribution of 5 nM Cas9 association and cleavage rates. Solid lines show two-fold changes (roughly the error of the assay). F) Summary of extent of cleavage across 61 sgRNAs from 0 to 20 bp of complementarity, relative to perfect target cleavage level. Cleavage level drops steeply with fewer than 17 complementary bases. G) Summary of extent of cleavage for other mismatch and bulge series across the length of the target.

Strikingly, virtually all perfect targets fell short of 100% cleavage (Figure 3C) even after incubating for an hour. While the dCas9 final fraction bound (f_final_) for perfect targets might be expected to fall significantly below 100%, the cleavage of template by active Cas9 over time would be expected to drive the reaction to completion in the limit of long incubations times. This sub-saturating behavior might be explained by Cas9 RNP binding to a target and, with some nonzero probability, entering a state where cleavage cannot occur and protecting the target. In general, the fraction of target bound exceeded the fraction cleaved (Figure 3C), supporting such a hypothesis. We also observed that, among perfect targets, final cleavage levels weakly correlated with final binding levels (R = 0.342, Figure 3D), suggesting that some of the variation in cleavage fraction may stem from variation in final binding levels, but the remainder is attributable to other sequence-dependent factors.

Other biochemical studies have concluded that cleavage is fast relative to rates of association (at 5 nM Cas9) for perfect targets (*12*, *13*). If cleavage were fast relative to association, we would expect a high degree of concordance between the observed association and cleavage rates for perfect targets because Cas9 association ought to be the rate-limiting step in both cases. We compared observed cleavage and association rates for perfect targets and found that cleavage rates were only moderately correlated with association rates (R = 0.465, Figure 3E). For many guides, we observe that perfect target cleavage is fast relative to association. However, a substantial fraction of guides induce cleavage more slowly than they associate, indicating that – for some guides – cleavage is indeed slower than Cas9 association (at 5 nM).

Previous work has shown cleavage is much more sensitive to imperfect matches than is binding (*22*) due to a conformational change required for target DNA cleavage (*7*, *27*, *28*). Our data are consistent with these findings. Across all sgRNAs, over 85% of targets with 17 bp of complementarity exhibited detectable cleavage (Figure 3F). Additional mismatches substantially decreased the fraction of targets cleaved: 38% of targets with 16 bp of complementarity exhibited cleavage below the threshold of detection, as did 62% of targets with 15 bp of complementarity (Figure 3F). In contrast, for most sgRNAs we observed only small changes in the final fraction bound for targets containing 15 bp of complementarity (Figure 2A).

The off-target cleavage data revealed an important trend: most targets with 15 or 16 bp of complementarity exhibited an intermediate level of final cleavage. In other words, off-target cleavage rates did not simply distribute near 0 (cleavage incompetent) and 1 (cleavage competent), but were instead broadly distributed (Figure 3G). The existence of a single mismatch or DNA bulge in any position had a modest impact on final cleavage levels. In addition to targets with less than 17 bp of complementarity, targets with RNA bulges of 2 or 3 nucleotides at positions −1 through −17, as well as targets with contiguous mismatches of 4 or more base pairs at any position, exhibited little cleavage despite high levels of final fraction bound (Supplementary Table 7).

### Target context modifies rate of Cas9 association and cleavage

In addition to assaying the 91 sublibraries described above, we also constructed two “3-mer scanning libraries” to test for the effects of flanking sequence on association and cleavage for λ1 and FANCF sgRNAs. These libraries were designed to harbor all possible trimers spanning the 5’ and 3’ flanks of the 23 bp target, extending 3 nucleotides 5’ (to position −23) and 6 nucleotides 3’ (to position 8) (Figure 4A). Binding data from this library revealed that only sequence variation near the 3’ end of the target site reliably produced large (>2-fold) changes in association rate of dCas9 (Figure 4B), and that association rates for targets with variation either 5’ of the target site or more than 3 bp from the NGG PAM rarely differed from the rate for the default flanking sequence (Figure 4B). While the typical association rate of Cas9 loaded with FANCF sgRNA was around a quarter that of a λ1 RNP, the effect of arbitrary 3’ nucleotides on the fold change of association rate relative to the default sequence context generally agreed, suggesting these context-specific effects on association are guide-independent (R = 0.775; Figure 4C; Supplementary Table 8). The identity of the base nearest the PAM was the most important feature governing cleavage rates, consistent with a previously reported NGGH motif for Cas9 (*29*). Relative Cas9 observed cleavage rates correlated with relative association rates (Figure S3; Supplementary Table 8), suggesting that at 5 nM Cas9, cleavage of perfect targets is fast relative to association for all flanking sequences for both tested sgRNAs. Thus, while flanking sequences clearly modify the rate of stable association, this assay lacks the time resolution necessary to assess effects downstream of association.

**Figure 4:**
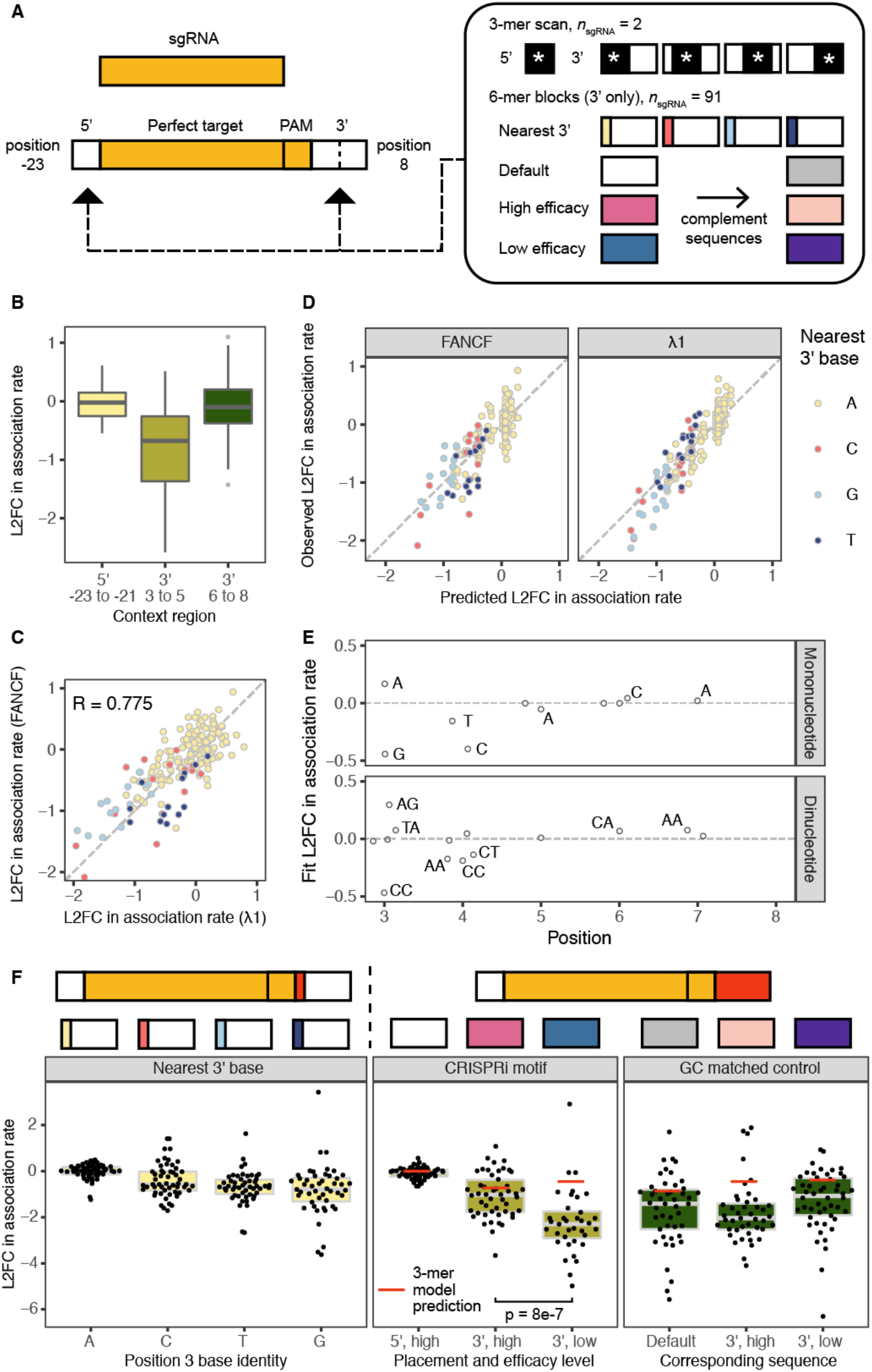
Impact of 5’ and 3’ sequence variation on perfect target binding and cleavage. A) 5’ and 3’ sequence variation was tested for all 3-mer blocks located next to or near the perfect target, each alternate base immediately downstream of the NGG PAM, and for specific 6-mer blocks 3’ of the perfect target. B) The impact of sequence variation 5’ of target sequence (positions −26 to −21) and 3’ of target sequence (positions 3 to 14) split by region (−26 to −21, 3 to 8, and 9 to 14). C) Comparison of 5’ and 3’ sequence variation effects from 3-mer scan for FANCF and λ1 sgRNAs. D) Results of learning a LASSO dinucleotide model for predicting the effect of 5’ and 3’ dinucleotides on association rate. E) Visualization of the selected coefficients from the LASSO dinucleotide model, distributed by position with high-weight features labeled. F) Effect of 3’ sequence variation on association rates across 91 RNPs. In left facet, the nucleotide immediately downstream of the NGG PAM modestly impacts on-rate. In center facet, extended motifs taken from a study of CRISPRi efficacy induce larger and more variable changes. In right facet, choosing GC-matched 3’ sequence demonstrates that motifs are not driven solely by GC-content.

We next attempted to model the impact of 6 bp of 3’ sequence variation on relative association rates for both FANCF and λ1. We converted these measurements of context effects into a matrix of dinucleotide features (104 total features) and measured log_2_-fold changes relative to the default perfect target sequence for each guide. An additive model fit by LASSO regression captured most of the variance (cross-validated R^2^ = 0.731, N = 411, Figure 4D). The fit parameters indicate that the presence of a G at the nearest 3’ position (NGGG extended PAM) slows association, in this case by 27% (Supplementary Table 9). However, as suggested by an analysis of CRISPRi/a data (*21*), an extended PAM consisting of a 3’ CC (NGGCC) slowed the association rate even more. When combined with an additional 3’ C (NGGCCC), the model predicted over a two-fold drop in association rate, more than double the reduction predicted for an NGGG extended PAM.

Context variants were also included in the 91 sublibraries to assess the guide-independence of effects of 3’ extended PAMs on association rate across a large number of guide sequences. To maintain library compactness, we tested the effects of five alternate 6 bp 3’ sequences and all three 1 bp substitutions downstream of the target on association to a perfect target sequence (Figure 4A). We chose the 6 bp blocks that exhibited the most (NGGCGGGAG) and least (NGG<u>GAATTT</u>) CRISPRi activity in Xu, et al (*20*) as well as complemented sequences to test if association preferences were driven by GC-content of the sequence blocks (Figure 4A).

Across context variants of all guides, association rates were typically the slowest to targets containing a G at the nearest 3’ base, consistent with an NGGH extended PAM motif for achieving the most rapid association (Figure 4F; Supplementary Table 10). While the median drop in association rate to an NGGG extended PAM was 1.7-fold, inserting the CRISPRi-disfavored sequence produced an even larger (6-fold) reduction (p = 8e-7 vs favorable). This effect is not only due to GC-content, as GC-matched controls showed smaller changes in their median relative association rates (2- to 4-fold decreases). The surprisingly large association rate decrease observed for this unfavorable 6-mer block was poorly predicted by the model trained on the scanning 3-mer data, suggesting that interactions beyond neighboring context nucleotides impact association rates. We also observed that association rates measured across perfect targets containing identical 6 bp blocks were more guide dependent and exhibited much larger variance than single base changes (1.8 vs 0.74 log_2_-fold units) (Figure 4F). These observations suggest that aspects of extended PAM preferences are guide dependent, and that while individual nucleotide changes have small effects, 6 or more nucleotide changes downstream of the PAM can lead to large differences in association rates for different sgRNAs.

### Concentration-independent mechanisms contribute to binding and cleavage specificity of Cas9

Our initial survey of the binding of Cas9 loaded with 90 different guide RNAs at 5 nM RNP affirmed two main points: the vast majority of library species exhibit intermediate levels of both binding (as measured by massively parallel filter binding) and cleavage. To determine if these behaviors can be described by a simple two-state binding model and to quantify the presence of nonproductive bound states, we selected 12 of the 90 guide RNAs for association profiling at 1.25 nM and 20 nM and cleavage profiling at 20 nM RNP.

Under a two-state binding model, the final fraction bound is a consequence of three independent parameters: protein concentration, k_on_ and k_off_. As protein concentration increases, the final fraction bound of a substrate also increases until it saturates at 100%. Yet, our extended dCas9 association data show that many Cas9 targets (e.g. a mismatch at position −4 for λ1, a mismatch at positions −4, −8, or −10 for ST3GAL5) do not saturate and instead plateau in their occupancy at levels far below 100%. Furthermore, for most sgRNA:target pairs, the observed final cleavage level was independent of Cas9 concentration (Figure S4A; Supplementary Table 11).

To address these discrepancies, we added an additional parameter to our fit capturing this ‘maximal productive binding’ to allow saturation below 100% of the DNA targets present in solution. Fitting the data in this manner thus models two phenomena: concentration-dependent initial binding affinity and concentration-independent entry into a stable non-canonical bound state. Our data were generally well fit by jointly fitting the three concentrations (Figure 5A; Supplementary Table 12), and these fits often returned maximal productive binding parameters well below 100%. We speculate that this sub-saturating binding behavior may be due to a bound state not detectable by nitrocellulose-mediated filter-binding, as documented previously for specific variants of LacR (*30*).

**Figure 5:**
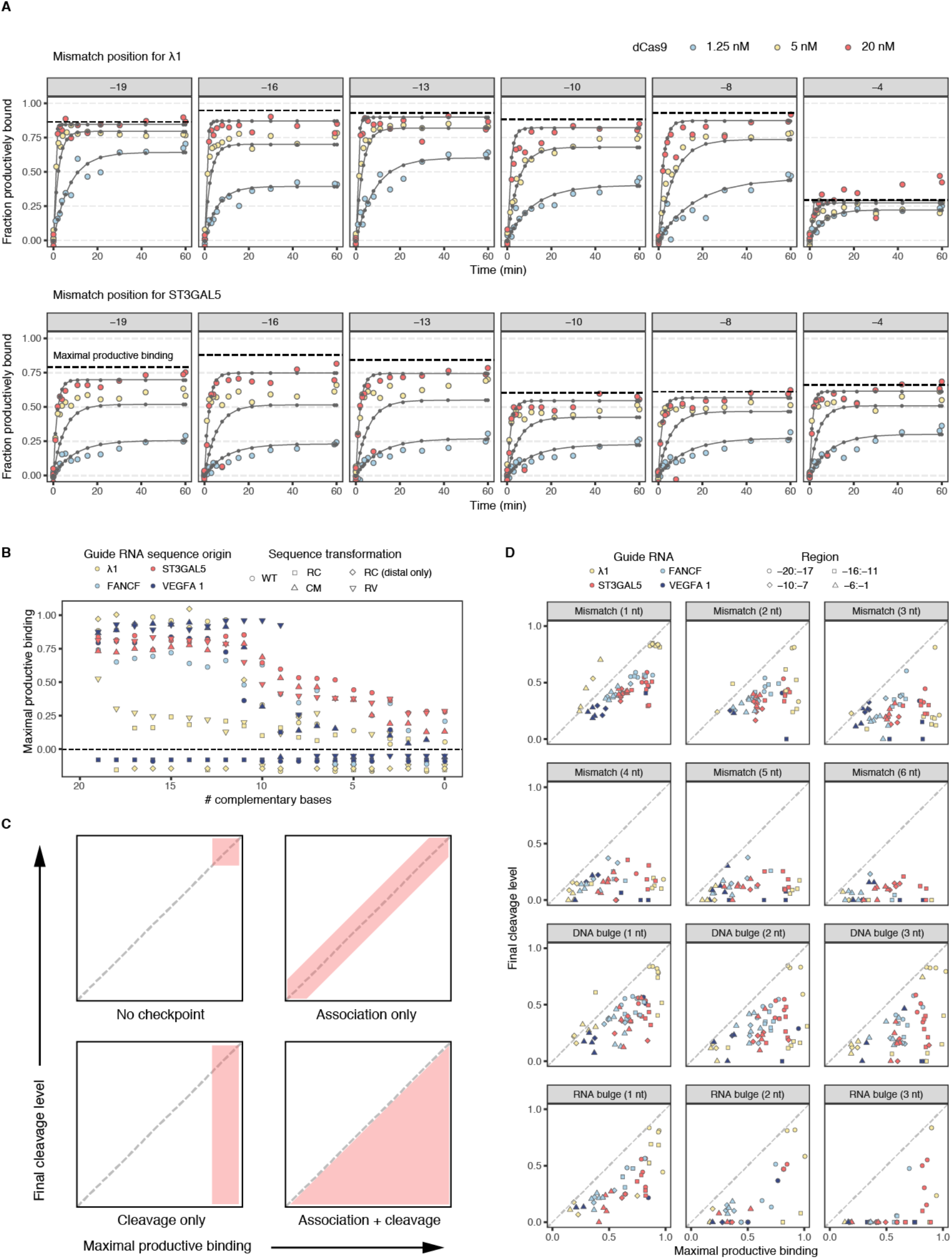
Joint fits for dCas9 association across three concentrations. A) Joint fits for two sgRNA and corresponding select singly mismatched targets. The dashed line signifies the fit maximal productive binding. B) Maximal productive binding is shown as a function of the number of complementary bases for all 12 jointly fit libraries. Most libraries show a transition around 10 base pairs of complementarity. Targets below the dashed line are below the limit of detection. RC: reverse complement sequence, CM: complement sequence, RV: reverse sequence. C) Expected joint distributions of maximal productive binding (x-axis) and final cleavage level (y-axis) under 4 different gating possibilities at Cas9 association and/or cleavage. D) Observed joints distributions for maximal productive binding (x-axis) and cleavage (y-axis) for jointly fit libraries. With increasing perturbation of sgRNA:target matching, targets fall farther below the diagonal.

Amongst the 12 guide RNAs we profiled, large differences in initial binding affinity (see methods) were observed only for off-targets of λ1 Cas9 RNP (and RNPs derived from λ1 sequence transformations) (Figure S4B; Supplementary Table 13). Unlike the other tested sequences, λ1-derived sequences are devoid of internal, non-canonical PAMs. It is likely that RNP-PAM interactions can dominate the initial binding affinity observed for targets with multiple PAMs. In these circumstances, most target mismatches may not change the kinetics of binding because initial binding is mediated by PAM interactions rather than target complementarity. Our maximal productive binding measurement instead appears to align with the conventional understanding of Cas9 specificity, exhibiting an 8-10 bp seed region that is sensitive to disruption, an 8-11 bp PAM-distal region that is largely resilient, and an intermediate zone sensitive to large perturbations (Figure 5B).

Because many targets appear incapable of attaining 100% productive binding, we hypothesize that there exist checkpoints in the binding process that can arrest Cas9 RNPs in nonproductive states. To understand the implications of our observations, we consider four possible models that either allow or disallow nonproductive states to trap target sequences prior to productive binding and/or cleavage. We explore the implications of these models under saturating protein concentrations ([Cas9]>>K_d_) (Figure 5C). In the simplest model, with no gating, Cas9 associates to a target in a single step and executes cleavage to completion. Under this model, all target sequences would cluster around 100% productive binding and cleavage. Under the second model, the addition of a cleavage checkpoint irreversibly halts or prevents cleavage of some targets, preventing 100% cleavage even with increased protein concentration or time. When we instead model a nonproductive, nitrocellulose binding-incompetent interaction, we expect identical sub-saturating behavior to arise in both association and cleavage data: RNP:target complexes forming nitrocellulose binding-incompetent interactions are prevented from progressing to cleavage, and all other targets are cleaved. Under our final model, gating occurs at both steps, such that final cleavage levels are bounded by the maximal productive binding level, which may in turn range from 0 to 100% (area below the diagonal in Figure 5C, lower right). This final model can produce two-fold sub-saturating behavior, at both binding and at cleavage stages.

We integrated the maximal productive binding estimates from our joint association fit data with the final cleavage level estimates from our 20 nM cleavage data to investigate the likelihood of each of the models described above. For a wide assortment of off-target sequences, the distribution of fit values strongly favors a model with sub-saturating behavior of Cas9 for both productive association and cleavage (Figure 5D). Targets with single RNA:DNA mismatches appear to exhibit extensive gating prior to productive binding, as measured by filter binding, but, of the fraction that appears bound, nearly all is able to cleave. Association and cleavage data for all other classes of off-target sequences are consistent with sub-saturating binding and cleavage. Interestingly, for each guide RNA, the extent of association gating is bimodally distributed. Off-targets with PAM-distal perturbations (positions −11 to −20) tend to cluster around 80% maximal productive binding, whereas targets with seed perturbations form a second cluster between 25 and 50% maximal productive binding. Saturation of cleavage, meanwhile, can range from 0 to 100%.

### Reversibility of Cas9 association declines over time

We previously noted that that longer RNP incubation times ultimately led to reduced dissociation (*15*). To characterize this phenomenon across diverse guides, we collected dissociation data series were after both 15 and 60 minutes of association with 20 nM dCas9 in a manner analogous to the association experiments (Figure 6A). Chase dsDNA without the sequencing adapters was used to quench dCas9 binding prior to adding samples to the nitrocellulose-covered vacuum manifold. λ1 targets with loss of PAM-distal complementarity demonstrated dissociation on the timescale of minutes (Figure 6B). As complementarity declined from 20 to 16 bp, the average off-rate for λ1 targets increased monotonically (Figure 6C; Supplementary Table 14). Our results were thus in line with our previous study of λ1 targets (*15*).

**Figure 6:**
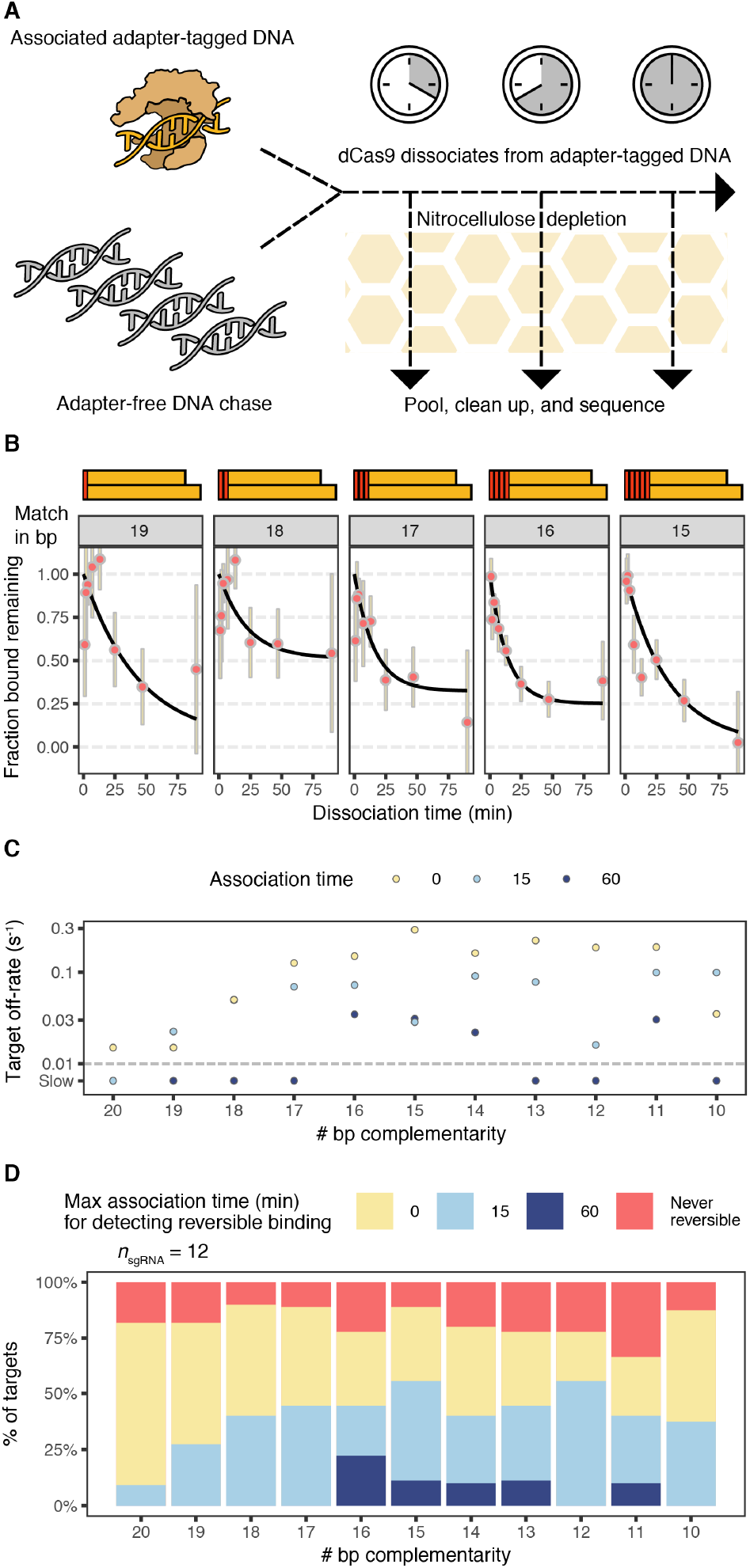
Quantification of dCas9 dissociation. A) Dissociation experiments rely on addition of an adapter-free DNA sink that blocks further binding to adapter-tagged libraries after initial variable-time incubation with 20 nM dCas9 RNP. B) Fraction bound to λ1 targets as a function of time and the number of PAM-distal mismatches. Fraction bound is normalized to that of the first timepoint after addition of the DNA sink. C) The observed λ1 sgRNA:dCas9 dissociation rate is shown as a function of the extent of complementarity (x-axis) and the time elapsed prior to the addition of DNA sink (point color). Yellow points (0 association time) are taken from the joint association fits across 3 dCas9 concentrations. Points below the dashed line are below limits of detection. D) Summary of observable dissociation across 12 sgRNAs. With greater association time, fewer sequences exhibit dissociation. Fraction colored red has fit off-rates below 0.01 per second in joint association fits.

Overall, we tested 6,865 off-target sequences across the same 12 sgRNAs measured across multiple dCas9 concentrations. Of these, 2,300 did not have sufficient binding prior to dissociation, 618 were fit to a negligible off-rate in the joint association fit, and 2548 exhibited dissociation below the limit of detection given the time scale for the dissociation experiments we conducted. The remaining 1,399 off-targets spread across the 12 guide RNAs, exhibited a similar pattern as seen for λ1: the loss of complementarity PAM-distally, from 20 bp to 16 bp, increased the observable dissociation, from 9% to 44% (for 15-minute association experiments) and from 0 to 22% (for 60-minute association experiments) (Figure 6D). This increase in the fraction of RNP:target complexes capable of releasing targets with PAM-distal mismatches supports the hypothesis that full target:guide pairing substantially reduces the reversibility of Cas9 binding.

### Cas9 binding and scission exhibit distinct sensitivities to target perturbation

Our results suggest a model wherein Cas9 traps off-target sequences in slowly acting or nonproductive states that both are not bound by nitrocellulose and block progression to cleavage (Figure 7A). Under this model, two concentration-independent parameters determine whether cleavage will occur at a target site when saturated with protein: the probability of productive binding and the probability of scission (conditioned on productive binding). The probability that a Cas9 RNP:target interaction cleaves an accessible target is the product of the two.

**Figure 7:**
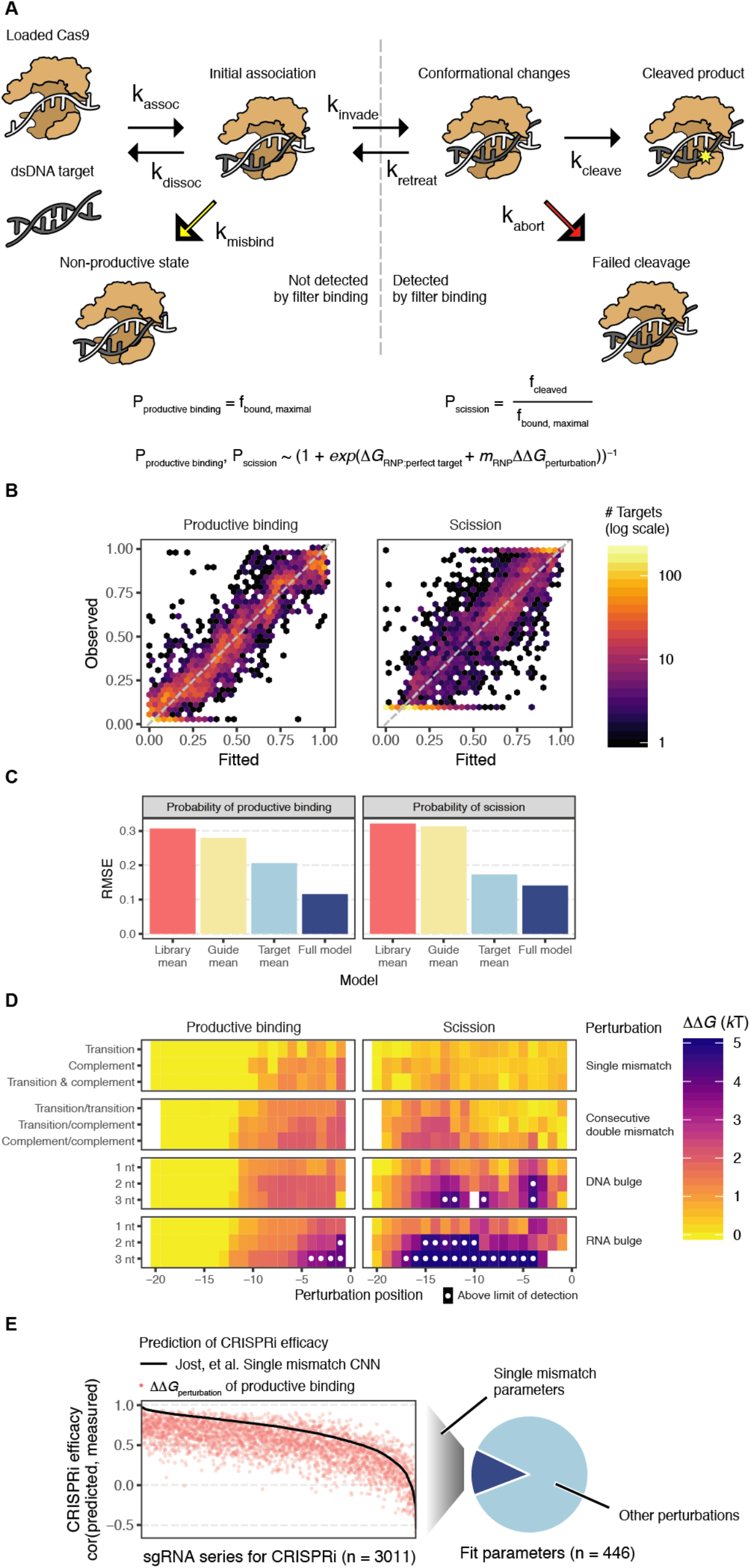
Model for Cas9 association and cleavage. A) Checkpoints exist both prior to productive association and successful nicking/cleavage. Reverse reactions from these nonproductive states are slow. The probability of productive binding (likelihood cleaved and aborted end states) is equal to the maximal productive binding estimate. The probability of scission (likelihood of cleaved state conditioned on productive binding) is equal to the final fraction cleaved divided by maximal productive binding. The full biophysical model for *P*_productive binding_ and *P*_scission_ as a function of *ΔΔG*_perturbation_, *ΔG*_RNP:perfect target_ and *m*_RNP_ is shown. B) 2D histogram of fitted and observed probability of productive binding and scission across all Cas9 RNPs. C) Root mean square error in predicting *P*_productive binding_ and *P*_scission_ of the full model compared to guessing the mean across all measurements or grouped by Cas9 RNP or target perturbation. D) Fit *ΔΔG*_perturbation_ values for select targets for productive binding and cleavage, plotted by perturbation position (x-axis) and type (y-axis). White circles represent measurements above the limit of detection. E) Results of comparing relative CRISPRi activity to productive binding *ΔΔG*_perturbation_ and Jost, et al. convolutional neural network predictions across sgRNA series, on left. Illustration of extent of target types with *ΔΔG*_perturbation_ values not scoreable by the Jost CNN, on right.

To learn about the sequence specificity of productive binding and scission, we designed one biophysical framework adaptable to both parameters. We first assigned each of the 12 Cas9 RNP:perfect targets pairs a baseline energy value to capture the partial productive binding and scission observed for perfect targets (*ΔG*_RNP:perfect target_). To group targets across Cas9 RNPs, we annotated mismatches (transition, complement, or both) and RNA and DNA bulges (from 1 to 3 nt) at each position for each RNP:target pair and defined targets with identical annotations as sharing the same “target perturbation” (Supplementary Tables 15-16). We then fit an energy penalty that decreases the likelihood of productive binding or scission (*ΔΔG*_perturbation_) to every target perturbation. Initial attempts at modeling suggested that different Cas9 RNPs exhibited differential sensitivity to sequence perturbations, and thus we also included an energy scaling parameter (*m*_RNP_) that allowed the overall magnitude of these energy perturbations to vary by guide.

Leave-one-out cross validation of RNP datasets suggested that perturbation penalties were stable and well-correlated with maximal productive binding estimates on held out data (mean spearman correlation of 0.81; Figure S5A). Using this framework, we fit productive binding energy penalties for 446 distinct target perturbations and both RNP-specific energy parameters for 11 dCas9 RNPs using 4,871 binding measurements, and, separately, scission energy penalties for 439 perturbations and RNP-specific energy parameters for 10 Cas9 RNPs using 3,603 binding and cleavage measurements (see methods; Figure 7B; Supplementary Tables 17-18). The logarithm of the productive binding energy scaling parameter was highly correlated with RNP:perfect target binding baseline energy (R = −0.78, p = 7e-3), as was the logarithm of the scission energy scaling parameter (R = −0.78, p = 8e-3; Figure S5B). From this we infer that more energetically favorable Cas9 RNP:perfect target pairs suffer commensurately larger penalties when mismatches disrupt their pairing, unexpectedly linking binding sensitivity to specificity.

Biophysical modeling demonstrated improved performance over taking the mean value per perturbation as measured by root mean square error, especially for productive binding (productive binding RMSE of 0.12 versus 0.21; scission RMSE of 0.14 versus 0.17; Figure 7C) and produced clear insight into how mispairing between sgRNAs and DNA targets influence productive binding and scission probability. Sequence perturbations at PAM-distal positions (−20 to −13) were universally assigned no energy penalty, and a 7-bp seed match was sufficient to discern some level of productive binding. Furthermore, productive binding loss due to PAM-proximal seed mismatches could be partially rescued by increasing complementarity PAM-distally, requiring approximately two additional distal matches to compensate for each seed mismatch (Figure 7C, Figure S5C). In contrast, scission is most perturbed by mismatches spanning positions −16 to −11 (Figure 7D), and a series of 6 or more mismatches anywhere between sgRNA and target usually abrogate scission activity (Figure S5C).

We also explored whether maximal productive binding energy penalties for doubly mismatched targets were additive with respect to their constitutive single mismatches. Consecutive double mismatches clearly diverged from additivity, in particular at PAM-distal positions that had no energy penalties for single mismatches exhibited energy penalties over 1 *k*T for double mismatches (Figure S5D; Supplementary Table 19). In contrast, nonconsecutive double mismatches that were at least 4 nucleotides apart appeared additive, suggesting that sufficiently distant mismatches may have independent effects on productive binding.

Finally, we evaluated whether productive binding *ΔΔG*_perturbation_ values could predict relative CRISPRi knockdown in human cells. Across 3,011 promoters with singly mismatched sgRNA series, we compared measured CRISPRi phenotypes both to our estimated *ΔΔG*_perturbation_ of productive binding and to activity predicted by a convolutional neural network (CNN) that incorporated additional features beyond the identity of the RNA-DNA mismatches (such as GC content and position relative to TSS) and was trained on this data set (Figure 7E) (*26*). The mean spearman correlation with measured CRISPRi activity was 0.508 for *ΔΔG*_perturbation_ versus 0.667 for the CNN. Thus, while a CNN specifically trained on these data outperformed our mismatch-only model, overall scores were remarkably similar (mean correlation between the models was 0.74), suggesting that biochemical parameters governing Cas9 binding and cleavage are the dominant features influencing in vivo efficacy. However, because the CNN model was trained only on single mismatch data, it is unable to predict more complex perturbations, whereas our *ΔΔG*_perturbation_ predictions span a broad variety of off-targets including 1-3 nucleotide bulges and mismatch series of arbitrary size that greatly expand the scope of off-target assessment. Surprisingly, 204 of the 386 more complex perturbations we estimate (53%) have predicted off-target activity within the range of predicted off-target activity for single mismatches, highlighting the vital importance of including such targets in in vivo off-target assessments.

## Discussion

Here we present a large corpus of *Streptococcus pyogenes* Cas9 binding and cleavage across diverse sgRNA sequences and corresponding DNA off-targets enabled by further parallelizing our pooled, sequencing-based filter binding assay. We now report an amortized cost of 8 cents per off-target measurement. In contrast to imaging-based methods that require maintaining fluidics and microscopes, our new design requires minimal equipment: principally a single 96-well vacuum manifold. We profiled ~10^3^ off-targets per RNP per experiment, and speculate that future applications of this technology to Cas9 or other DNA- or RNA-binding proteins of interest could assess more than 250,000 targets with straightforward protocol modifications. Thus, we believe massively parallel filter binding represents cost-effective and operationally straightforward tool for profiling protein-nucleic acid binding kinetics.

In this study we show that differences in perfect target association kinetics appear to explain some of the differences in screening efficacy across sgRNAs. Two underappreciated phenomena – sgRNA folding and disadvantageous extended PAM sequences – appear to modify efficacy at the level of binding, with implications for both CRISPRi/a and CRISPR KO screens (*25*). To investigate this possibility, we compared perfect target biophysical measurements to CRISPRia scores for sgRNAs with CRISPRi measurements (*21*) and found that empirical measurements of RNP association exhibited greater predictive power for sgRNA efficacy. As a filter-binding experiment is generally simpler and faster than a CRISPRi-based screen in cells, we believe that measurement of association rates *in vitro* may be a useful alternative to computational and cell-based methods for evaluating guide efficacy.

Yet, altered Cas9 RNP association to off-targets does not appear to explain reduced activity at off-target sites. We observed that association kinetics of off-target sequences usually cluster tightly around that of their respective perfect targets. Thus, changes in off-target activity are unlikely to be governed by differences in k_cat_ or k_on_, consistent with suggestions of the mechanism of high-affinity binding by other nucleic-acid guided proteins (*31*). Reducing off-target activity by increasing off-rates via PAM distal mismatches (*15*) has some utility, as RNP:off-target complexes exhibited less dissociation over time. Furthermore, initial binding affinity suggested that little more than a few PAMs might be adequate for appreciable dCas9 occupancy. Both observations appear in conflict with the high reported specificity of sgRNAs in CRISPRi screens.

We also observe that scission of many bound DNA targets is incomplete, supporting a branching rather than linear (*28*) binding and cleavage process involving intermediate states. Other investigators have attributed incomplete cleavage to the existence of a nonproductive state comprising 15% of the RNP:target complex and slow biphasic reaction steps in Cas9 catalysis (*10*, *12*, *13*). The expanded scope of our off-target dataset strongly suggests that the probability of cleavage varies both by sgRNA sequence and the number and type of DNA target mismatches present. The most likely explanation is that the probabilities of both stable binding and scission of stably bound targets are strongly dependent on the sequence identity of the RNP:target pair in a concentration-independent manner, ranging both above and below the 15% seen for commonly studied RNP:target pairs. This behavior suggests that multiple checkpoints have evolved to mitigate Spy Cas9 off-target activity independent of Cas9 RNP:target interaction affinity. Unexpectedly, we show that Cas9 filter-binding experiments appear to reflect additional state information beyond the binary notion of bound or unbound. Specifically, off-targets with numerous sgRNA:target mismatches are rarely fully depleted when passed through nitrocellulose, and increasing RNP concentration does not enhance depletion. The inferred maximal depletion for an RNP:target pair consistently serves as an upper bound for the fraction of target that can be cleaved, suggesting this state is also not cleavage competent. We speculate that off-target sites trap Cas9 in slowly acting or nonproductive states that disassemble when passed through nitrocellulose (Figure 7A).

The manner by which Cas9 engages with on- and off-target sites has clear practical relevance for application of Cas9 technologies. A model of dCas9 binding similar to ours has been proposed to explain how mismatched sgRNAs can permit concentration-independent, “noiseless” CRISPRi-mediated gene silencing in bacteria (*32*). The authors observe that dCas9 resists eviction by RNA polymerase extension and blocks gene expression with a fixed probability, *P*(stop), that positively correlates with target complementarity. We speculate that *P*(stop) may be functionally equivalent to what we report as the probability of productive binding, and that nonproductive Cas9 RNP:target interactions are easily dismantled by either colliding with RNA polymerase or passing through nitrocelluose. Thus our work adds to the growing biochemical evidence of nonproductive bound states (*12*, *13*, *33*).

Understanding off-target association and cleavage may prove key to engineering workhorse variants of CRISPR enzymes. Most studies have focused on optimizing Spy Cas9 cleavage (*9*, *34*, *35*), and a recent preprint confirmed that the association kinetics for the most widely-used engineered Cas9s do not differ from their wild type counterparts (*36*). Yet, engineering efforts designed around Spy Cas9 binding have achieved greater on-target efficacy and specificity (*37*). More broadly, off-target detection methods have demonstrated substantial time- and concentration-dependent off-target activity (*38*, *39*). For this reason, protein engineering efforts are unlikely to offer a single solution for experiments that operate over different time scales with different tolerance for off-target effects, and more advanced biophysical models for Cas9 activity remain a top priority.

Despite the efforts of several groups, predicting the kinetics and thermodynamics of binding for an arbitrary RNP complex to its perfect target remains an outstanding challenge. Indeed, new 3’ sequence requirements for Cas9 binding were only recently discovered (*40*), which we confirm (Figure S6). While initial RNA-seq data showed little to no off-target activity of CRISPRi (*41*), new results from screens of noncoding elements in human cell lines (*42*) and screens of essential genes in bacteria (*43*, *44*) suggest that a variety of sequences remain difficult to target without the possibility of substantial off-target effects. Interestingly, we observe that guide sequences exhibiting strong on-target binding typically have more specific binding behavior, with implications for rectifying poor gRNA performance. We anticipate that the generation of large-scale data on off-target binding, as well as detailed thermodynamic modeling of potential binding and cleavage events, will become only more important as an increasing number of guide sequences are deployed for therapeutic applications.

## Materials and Methods

### (d) Cas9 RNP preparation

sgRNAs were in vitro transcribed using the NEB EnGen sgRNA synthesis kit (catalog #E3322S) according to manufacturer’s instructions, starting with 0.15 reaction units per sgRNA and scaled up to 0.5 units as needed to generate sufficient material for each sgRNA. sgRNA were purified using Agencourt RNAClean XP beads for the all-sgRNA round (part #A63987) and Zymo RNA Clean & Concentrator-5 (catalog #R1013) for additional syntheses. Cas9 and dCas9 was provided by the Doudna lab.

For loading, each sgRNA was incubated at 98C for one minute and slowly cooled to room temperature. dCas9 was diluted to 100 nM and incubated with an equal volume of sgRNA at 20% excess in 1X binding buffer (20 mM Tris·HCl, pH 7.5, 100 mM KCl, 5 mM MgCl_2_, 5% glycerol, 0.05 mg/mL heparin, 1 mM DTT, and 0.005% Tween 20), a final working concentration of 50 nM. Loaded dCas9 is further diluted to attain the desired concentration (1.25 nM, 5 nM and 20 nM) for association experiments and 20 nM for dissociation experiments in 1X binding buffer.

### Library preparation

Guide RNAs were curated from a variety of sources, including genetic screens, Cas9 off-target screens, efforts to characterize Cas9 biochemistry, and, lastly, sequence transformations of the above. Sequence transformations consisted of taking the complement, reverse, or reverse complement of parts of guide RNA sequences in order to allow direct comparison between nucleotide composition-matched guide RNA sequences. Across all off-target types, 46,393 off-target sequences were designed, each for one of the 90 curated sgRNAs (plus duplicate sublibrary for the λ1 sgRNA).

In addition to the 23-bp target and 6-bp 5’ and 3’ flanking sequence contexts, each sublibrary of on-target and corresponding off-target sequences was assigned a 13-bp handle for separate amplification (*45*) filtered to be free of GG and CC dinucleotides plus 17- and 18-bp universal adapters on either end of the target to enable amplification of the entire pool. Oligonucleotides were synthesized in a single pool by CustomArray on a 92,918 array (each sequence in duplicate) and PCR amplified using NEBNext 2X master mix (catalog #M0541L).

Following the initial amplification, each sublibrary was amplified with 16 distinct pairs of barcoded forward and reverse primers in separate reactions (98 C denaturation, 68 C annealing, 72 C extension). PCR products were purified using Ampure beads and quantified by Qubit dsDNA HS kit (catalog # Q32854) prior to dilution to 1 nM total oligo working concentration.

### Massively parallel filter-binding experiments

Custom-designed adaptors for loading samples into a vacuum manifold were ordered online via 3D printing from 3D Hubs (ABS FDM, 40% infill, 200 um resolution). Adaptor surfaces were sanded down with 300-grit sandpaper to remove striations left by 3D printing. Prior to use, surfaces were coated with a superhydrophobic residue to prevent sample loss to wetting of the surface. Rust-Oleum NeverWet (Amazon) Step 1 was first applied in 2-4 short bursts of spraying and dried in a fume hood for 2 hours. One coat of Step 2 was applied and then left to dry overnight. A second coating was applied the next day and fully dried prior to use. Hydrophobic residues remained intact for a week but deteriorated and required new coatings for peak performance if left for longer periods.

The filter-binding vacuum manifold systems was assembled by inserting a sterile 1mL deep 96-well plate into the bottom of a 96-well vacuum manifold, placing the upper half over the plate, layering a cut section of Fibre Craft foam (Amazon) over the surface of the plate and adding the custom adaptor to reach into the wells. To prepare nitrocellulose, a precut membrane was soaked in binding buffer before transferring to the surface of the adaptor to create a vacuum-tight seal.

For association experiments 1.25 nM, 5 nM, and 20 nM dCas9 (10 nM dCas9 for context experiments) was incubated with 16 barcoded libraries individually (final library concentration 100 pM) in 40 μL of 1X binding buffer at room temperature, timed to yield measurements at 1, 2, 3, 4, 5.5, 8, 11, 15.5, 21.5, 30, 42, 59 and 60 minutes of association plus three zero timepoints. For dissociation experiments 20 nM dCas9 was incubated with 14 barcoded libraries at room temperature for variable association time followed by addition of a final concentration of 40 nM competitor on-target DNA to yield measurements at 1, 2, 3.5, 7, 13, 25, 47, and 90 minutes of dissociation plus two pre-dissociation samples and four zero timepoints.

Each association and dissociation timepoint reaction was passed through the nitrocellulose filter and flow-through collected from the corresponding wells. Samples for 6 sublibraries were pooled and purified using Qiagen MinElute columns. Libraries were quantified by Qubit dsDNA HS assay in the case of association experiments and by qPCR with a standard curve derived from a Qubit-quantified dsDNA library in the case of dissociation and cleavage experiments. All libraries were sequenced PCR-free using Illumina Nextseq v3 chemistry with 2×75 reads.

### Cleavage experiments

Loaded active Cas9 is added to barcoded target libraries in binding buffer followed by quenching with 16 mM EDTA and placing on ice, timed as in the association experiments. Following EDTA quench, reactions were immediately incubated at 65 C for 10 minutes to deactivate Cas9. Reactions were pooled and cleaned up using Qiagen MinElute columns as above.

### EMSA experiments

DNA oligos were ordered from IDT such that a forward oligo with 6 base pairs of sequence upstream of the Cas9 target sequence partially overlapped reverse oligos with variable numbers of bases downstream of the target (7, 16, or 20 bp). A reverse, Atto532-labeled oligo that extended 20 bases downstream was ordered in parallel to permit visualization of results on a Typhoon imager. All reverse oligos were annealed and extended with the forward oligo using NEBNext 2X master mix. Labeled DNA was added to dCas9 RNP with or without unlabeled competitor DNA of uniform length present (again either 7, 16, or 20 bp downstream of the target), for final concentrations of 200 pM labeled DNA, 5 nM dCas9 RNP and 20 nM competitor. Bound and unbound labeled DNA was separated by electrophoresis using Novex 10% TBE precast gels (catalog # EC6275BOX).

### Sequence read data analysis

Fastqs were first trimmed of adapters using SeqPurge (*46*). Trimmed forward and reverse reads were merged using FLASH (*47*) with the max-mismatch-density parameter set to 0.01 and the min-overlap parameter set to 10. Merged fastq reads were assigned to target library sequences permitting one single nucleotide mismatch in the sublibrary primer sequence and an exact match throughout the rest of the target. Reads were aggregated by target to produce a count table of counts per target and timepoint.

### Estimating final fraction bound, final fraction cut, initial binding affinity and maximal productive binding

For single concentration associations, count data for each target was fit to the following equation in R using the `nls` function:

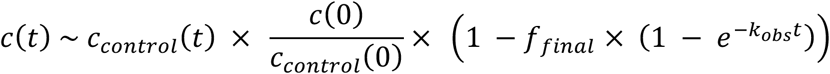

c(*t*) is the target sequence count at timepoint *t*. c_control_(*t*) is the control sequence count at timepoint *t*. f_final_ is the final fraction bound, and k_obs_ is the observed rate constant. f_final_ was initialized to 0.9 and k_obs_ to 0.024 per nM per minute times the Cas9 concentration. The control parameter was set to `nls.control(maxiter=300,warnOnly=TRUE)`.

For data visualization, fraction bound [f_bound_(t)] was inferred as follows:

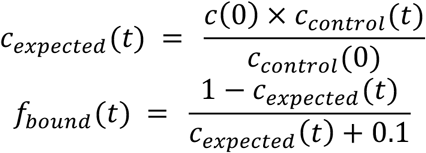

0.1 was added to the denominator to prevent divide-by-zero errors in the rare case of zero reads at timepoint 0. 90% confidence intervals for inferred fraction bound were calculated by adding and subtracting 1.64 times the square root of c(*t*) for each timepoint and calculating final fraction bound as before.

Some targets were not fit due to the following criteria:

1. Binding was not dynamic over the course of the experiment. The average final fraction bound at the two latest timepoint did not exceed the upper limit of the 90% confidence interval for each of the first two timepoints.
2. Counts for the target sequence were too small to be reliable. There were fewer than 5 timepoints in the first half of the experiment that exceeded 30 reads.

Targets that did not meet both of the above requirements were further stratified. Of the targets that were not dynamic, those that had at least 5 timepoints in the second half of the experiment with the lower limit of the 90% confidence interval above 0 were fit to a horizontal line based on the timepoints in the second half of the experiment (no rate parameter). Of the remaining targets, those that averaged below 15% final fraction bound in the second half of the experiment were flagged as low affinity. The remainder (i.e. targets with large confidence intervals and small change in final fraction bound from the beginning to the end of the experiment) were annotated as noisy.

We observed that after performing initial fits, some timepoints were consistent outliers across DNA targets in the same experiment. This could be explained by a biased control target count for such outlier points, which would affect the inference of fraction bound for all other targets. To address this, timepoints for which the magnitude of the averaged residuals exceeded 2.5 times the median magnitude of the averaged residuals were excluded, and count data was refit using the remaining timepoints. The association rates we report refer to f_final_ times k_obs_. Cleavage data were fit in the same manner as binding data.

For joint association analysis across three dCas9 concentrations, measurements outside an inferred fraction bound from 0 to 150% were excluded as outliers. After filtering, the following equation was used:

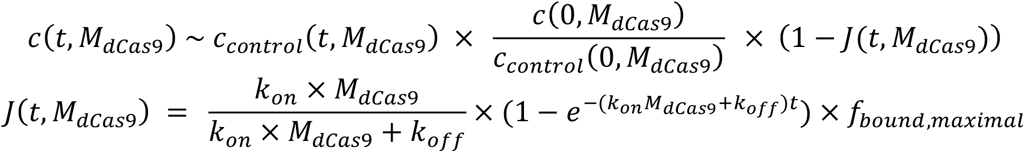

M_dCas9_ is the concentration (M) of dCas9. f_bound,maximal_ is the maximal productive binding.

For joint fits, k_on_ was initialized to 2e7 per M per minute, k_off_ to 0.02 per minute, and f_bound,maximal_ to 0.85.

Initial binding affinity was calculated from k_on_ and k_off_ in the manner of a K_d_, reported in units of *k*T:

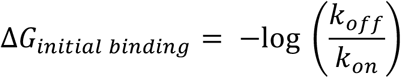

As in the association experiments, targets in the dissociation experiments were required to exhibit a 15% drop in fraction bound over the course of the experiment to qualify (to be sufficiently dynamic). Sequences that did not start above 15% bound prior to beginning dissociation were deemed low affinity and removed from consideration. Only timepoints following quench (*t* > 0) were included in the fit.

Dissociation experiments were fit to a distinct equation:

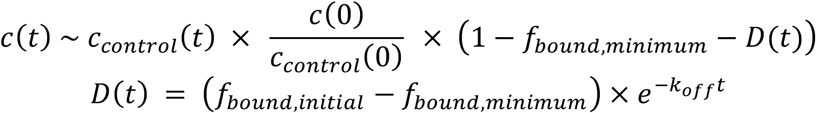

f_bound,minimum_ is the fit fraction of dCas9 that did not reverse on the timescale of the experiment. f_bound,initial_ is the fit fraction of dCas9 bound at *t* = 0.

After fitting curves, fits with an k_obs_ or k_off_ below 0.02 per minute, an observed rate above 2 per minute, an f_final_ above 1.2 or an f_final_ below −0.2 were excluded as poor fits.

The LASSO model for 3’ context effects was fit using the R package glmnet. Coefficients for dimer identities by position were retrieved by running the `coef` command with parameter `s = “lambda.1se”`.

### Defining biophysical model parameters for productive binding and scission probabilities

We assume that the choice for a specific Cas9 RNP:perfect target pair between entering a productively bound state or a nonproductively bound state can be modeling with a simple energy gap: *ΔG*_RNP:perfect target_, whereby more negative energies favor productive binding. We anticipate that most Cas9 RNPs should have a value near or below zero, such that that the probability of productive binding is near or above 50%. In addition, we assign each sequence perturbation a fixed adjustment to the energy gap that applies independent of Cas9 RNP: *ΔΔG*_perturbation_. However, it stands to reason that different Cas9 RNP:target pairings will be differentially impacted by mismatches and bulges. Specifically, it is expected that RNP:target pairs that are more energetically favorable should conversely suffer larger energy penalties when disrupted. We first attempted a parameter-free correction by using DNA:DNA and RNA:DNA duplex hybridization energies estimated by MELTING5, but performance was poor. Instead, we introduced another parameter (*m*_RNP_) to scale *ΔΔG*_perturbation_s per RNP.

From these parameters, we derived the probability of productive binding:

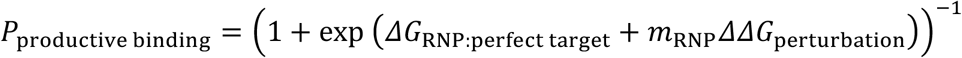

The same equation was used for probability of scission.

### Fitting biophysical model parameters for productive binding and scission

Maximal productive binding data was organized into a matrix of sequence perturbations by dCas9 RNP guide sequences. Maximal productive binding levels were subject to a series of quality control steps:

1. Overly large estimates of maximal productive binding (> 150%) were replaced with NAs
2. Alternate perfect target contexts were removed to ensure one value for *ΔG*_RNP:perfect target_
3. Targets with slow initial k_on_ (< 2000000 M^−1^ min^−1^) or fast initial k_off_ (> 1 min^−1^) estimates were replaced with 2% maximal productive binding
4. Targets with high estimates of maximal productive binding (> 98%) were replaced with 98%.
5. Targets with low estimates of maximal productive binding (< 2%) were replaced with 2%.
6. Sequence perturbations with fewer than 4 valid maximal productive binding estimates (from different RNPs) were removed

After filtering, 465 perturbations remained for fitting, with 19 redundantly encoded perturbations. One RNP (the reverse complement of VEGFA site 1) was excluded entirely due to low levels of productive binding and few valid fitted values, leaving 11 columns for a 465 by 11 data matrix *d*.

All productive binding parameters were fit jointly using the `nls.lm` function in R. The geometric mean of the *m*_RNP_ parameters was constrained to 1 by fitting only 10 free parameters and inferring the 11^th^. *m*_RNP_s were initialized to 1, and *ΔG*_RNP:perfect target_ values were initialized to 0. *ΔΔG*_perturbation_ values were initialized by converting the difference in probability of productive binding from the perfect target to the perturbed target to a *ΔG* and taking the average across all dCas9 RNPs. If the average for a perturbation was below −0.1 *k*T, it was set to −0.1 *k*T.

*m*_RNP_ values were bounded between 0.2 and 5. *ΔG*_RNP:perfect target_ values were bounded between −6 and 3 *k*T. *ΔΔG*_perturbation_ values were bounded between −0.1 and 6 kT.

The data matrix m was predicted as follows:

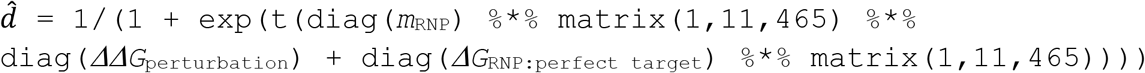

Residuals were reported to `nls.lm` by taking the difference between 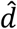 and *d*, removing NA values, and converting the matrix to a vector. After fitting, *ΔΔG*_perturbation_s below 0 were set to 0.

After an initial fit, the top 20 perturbations ranked by mean absolute deviation were manually examined for potential outliers. Out of the 220 measurements examined, 9 dCas9 RNP:off-target pairs appeared to have extreme values and were designated as outliers. In a majority of cases, refitting the data had a marginal impact on the fitted values, suggesting that overall, fits were robust to random error.

Leave-one-out cross validation was performed by removing one column from *d* and taking the Spearman correlation between learned *ΔΔG*_perturbation_ values and estimated probabilities of productive binding (which would be unaffected by *ΔG*_RNP:perfect target_ or *m*_RNP_).

Probability of scission data was fit in much the same way, with some added steps and modifications. Measurements where the probability of productive binding was below 2% were replaced with NAs. Final cleavage levels above 99% were replaced with 99%. The probability of scission was calculated as maximal productive binding divided by final cleavage level. Probabilities of scission above 99% were replaced with 99%. Probabilities of scission below 10% were replaced with 10% because low levels of scission were difficult to resolve, especially when the probability of productive binding was low.

In total, 458 perturbations were able to be fitted, although one additional Cas9 RNP (reverse complement of the distal sequence of λ1) was removed because of low levels of cleavage across all targets. *ΔΔG*_perturbation_ values were bounded between −0.1 and 7 kT because probability of scission for perfect targets generally exceeded the probability of productive binding for perfect targets which increased the range of detection from *ΔG*_RNP:perfect target_ values. Finally, out of the 200 measurements for the 20 perturbations with greatest error, only 4 measurements were deemed outliers.

## Supporting information

All supplementary tables

## Contributions

E.A.B. conceived of and led the study, with guidance from W.J.G. E.A.B., H.B.B. and W.R.B. developed filter-binding methodology and performed experiments. J.S.C. and J.A.D. provided protein. W.J.G supervised the study. All authors contributed to writing the manuscript.

## Data and materials availability

Raw count data and 3D printing design file are available on Figshare: https://doi.org/10.6084/m9.figshare.12526157.v1

## Supplementary Materials

### Supplementary tables

ST1: Oligo and target sequences for the primary (d)Cas9 binding and cleavage sublibraries.

ST2: Counts of target types by sublibrary and filtering criterion for analysis.

ST3: Rule Set 2, CRISPRia and association rates for effective and ineffective CRISPRi guide RNAs profiled in this study

ST4: Results for 5 nM dCas9 association profiling across all sublibraries

ST5: Annotated DNA bulge association data

ST6: Predicted guide RNA activities from Jost, 2020 for dCas9 binding targets

ST7: Results for 5 nM dCas9 cleavage across all sublibraries

ST8: 10 nM dCas9 binding and cleavage results for FANCF and λ1 sublibraries

ST9: LASSO coefficients for the effects of context variation on association rate

ST10: Association rate changes for context variants across all sublibraries

ST11: Results for 20 nM dCas9 cleavage for the chosen sublibraries

ST12: Joint fits for 1.25, 5, and 20 nM dCas9 association data

ST13: Initial k_D_ values estimated from joint fit results

ST14: Results for dCas9 dissociation profiling of the chosen sublibraries

ST15: Probability of productive binding for each target perturbation and RNP

ST16: Probability of scission for each target perturbation and RNP

ST17: *ΔΔG*_perturbation_ values for productive binding and cleavage

ST18: Results of cross validation on *ΔΔG*_perturbation_ estimates

ST19: Double mismatch *ΔΔG*_perturbation_ values for productive binding alongside constitutive single mismatches

### Supplementary figures

**Figure S1:**
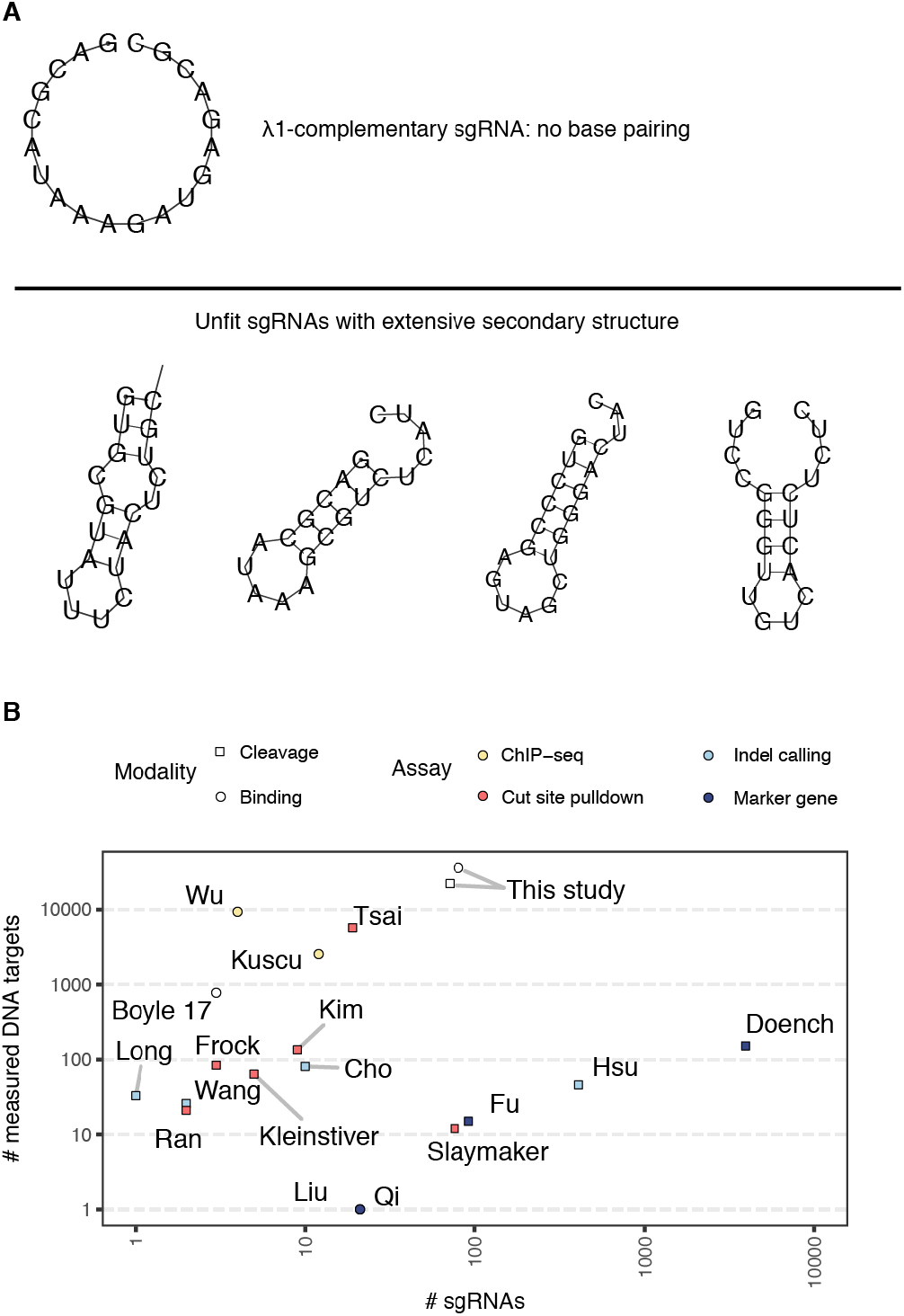
A) For most programmed targets, RNAfold predicted little secondary structure outside of the intended sgRNA hairpins (above). Amongst sgRNAs with otherwise inexplicably poor activity, extensive secondary structure was common (below). B) The number of DNA off-targets profiled contained in the present study exceeds that of other recently published datasets as aggregated by Zhang, et al. While other technologies have larger scale for assessing sgRNA cleavage activity, filter-binding permits profiling of a larger number of sgRNAs for binding.

**Figure S2:**
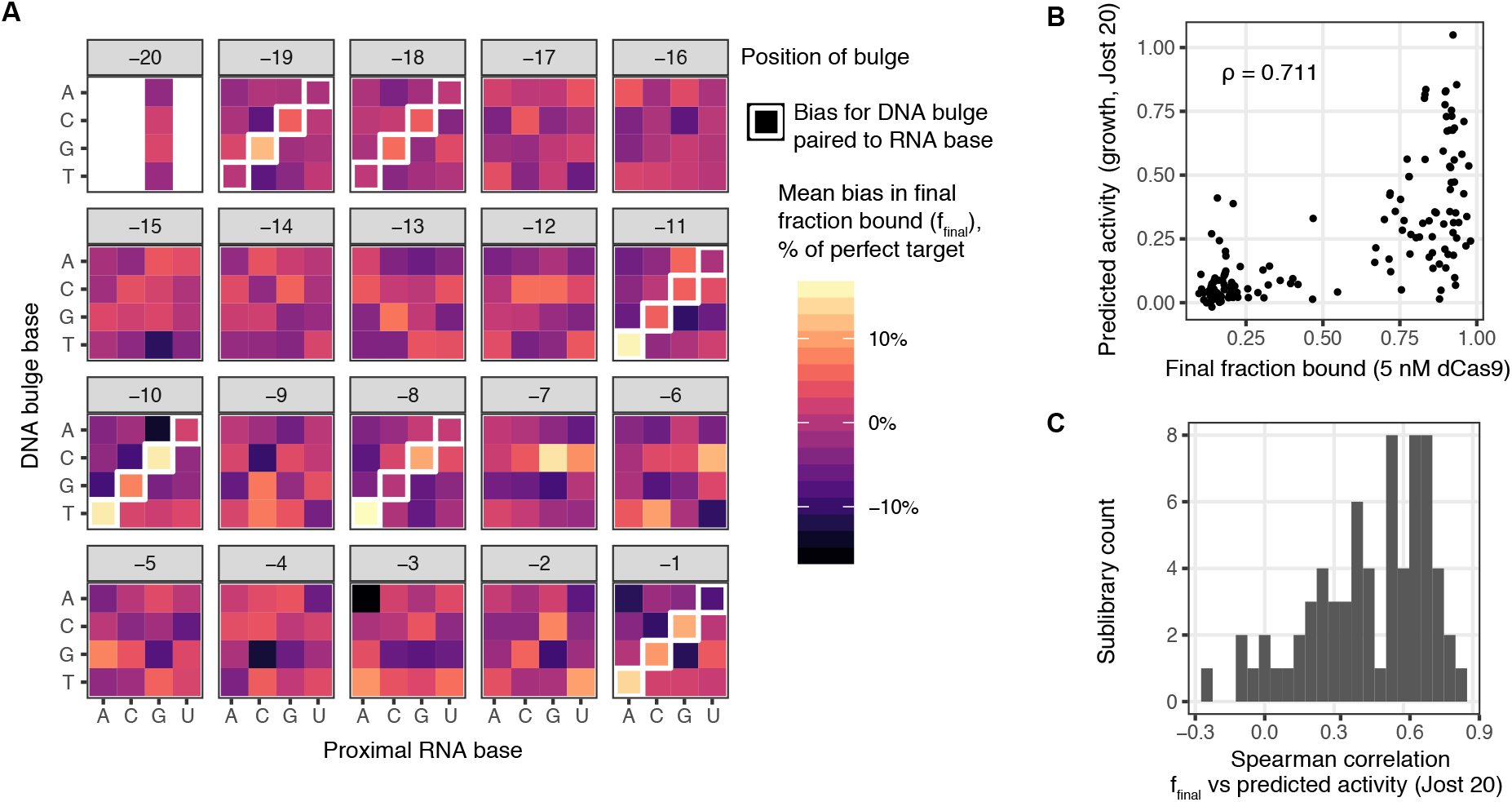
A) For each sgRNA position and reference DNA base, the difference between f_final_ for a specific DNA bulge and the mean f_final_ of all 4 bulges is shown. B) Plotting off-target f_final_ against predicted CRISRPi activity for the λ1 sublibrary reveals substantial binary agreement between the two measures. C) A histogram of the Spearman correlations between f_final_ and predicted CRISPRi activity per sublibrary shows predominately positive correlations.

**Figure S3:**
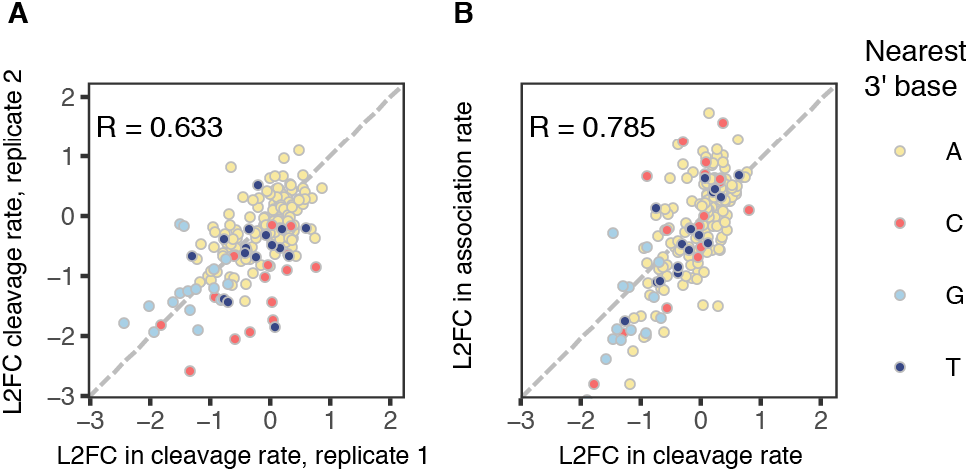
Plotting log2-fold changes in cleavage rate for perfect targets relative to the default target vs A) replicate measurements and B) log2-fold changes in association rate shows how sequence variation 5’ and 3’ of the target site influences association, which cascades to cleavage kinetic differences.

**Figure S4:**
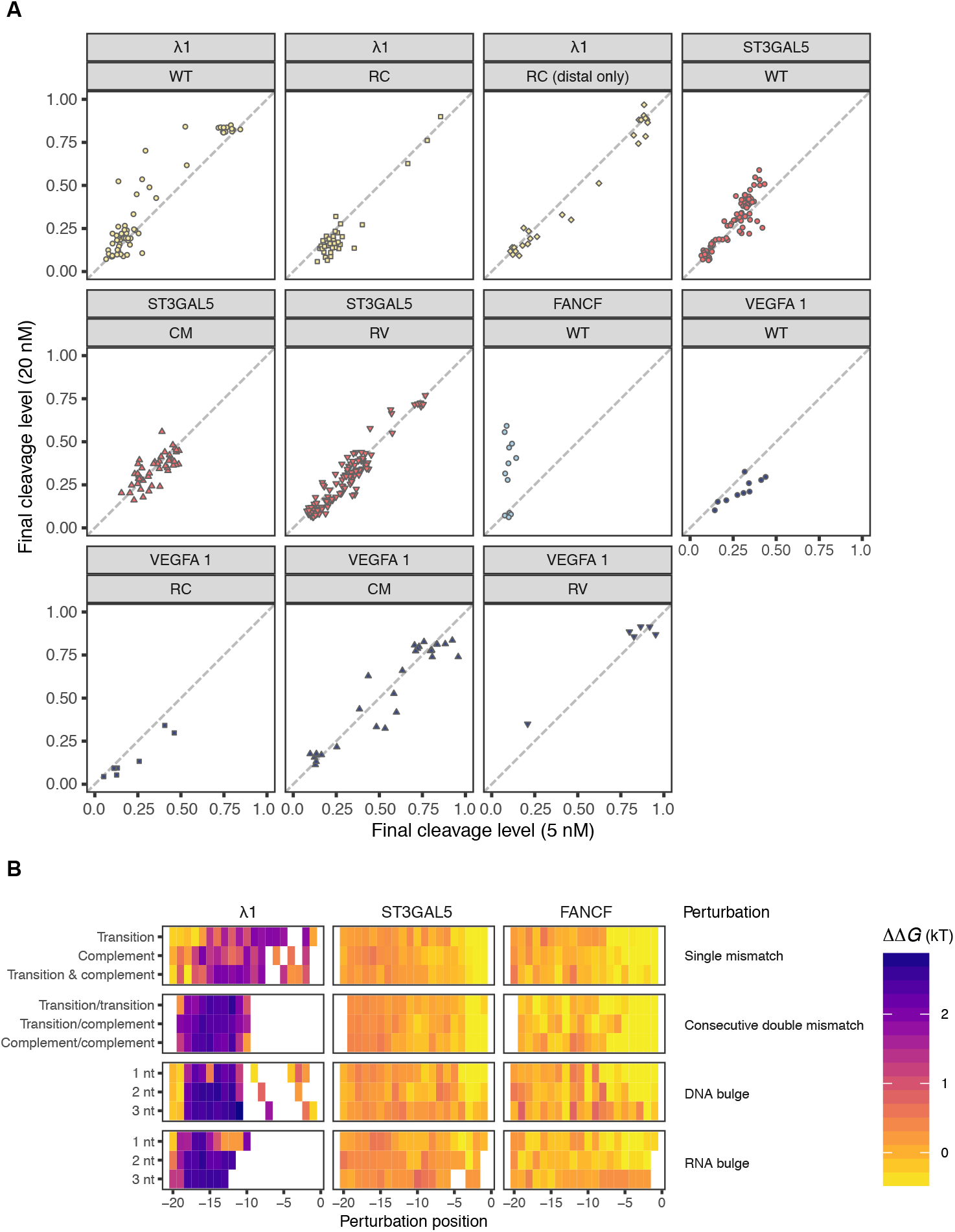
A) Pairwise comparison of final cleavage levels for 11 Cas9 RNPs and corresponding sublibraries for 5 nM and 20 nM Cas9. Mismatch series are shown for each sublibrary. The diagonal represents equal final cleavage level. B) Initial binding affinity as measured from initial k_D_ estimates from jointly fit dCas9 RNP association experiments, relative to their respective perfect targets. Larger *ΔΔG*s imply an increase in initial off-rates or a decrease in initial on-rate.

**Figure S5:**
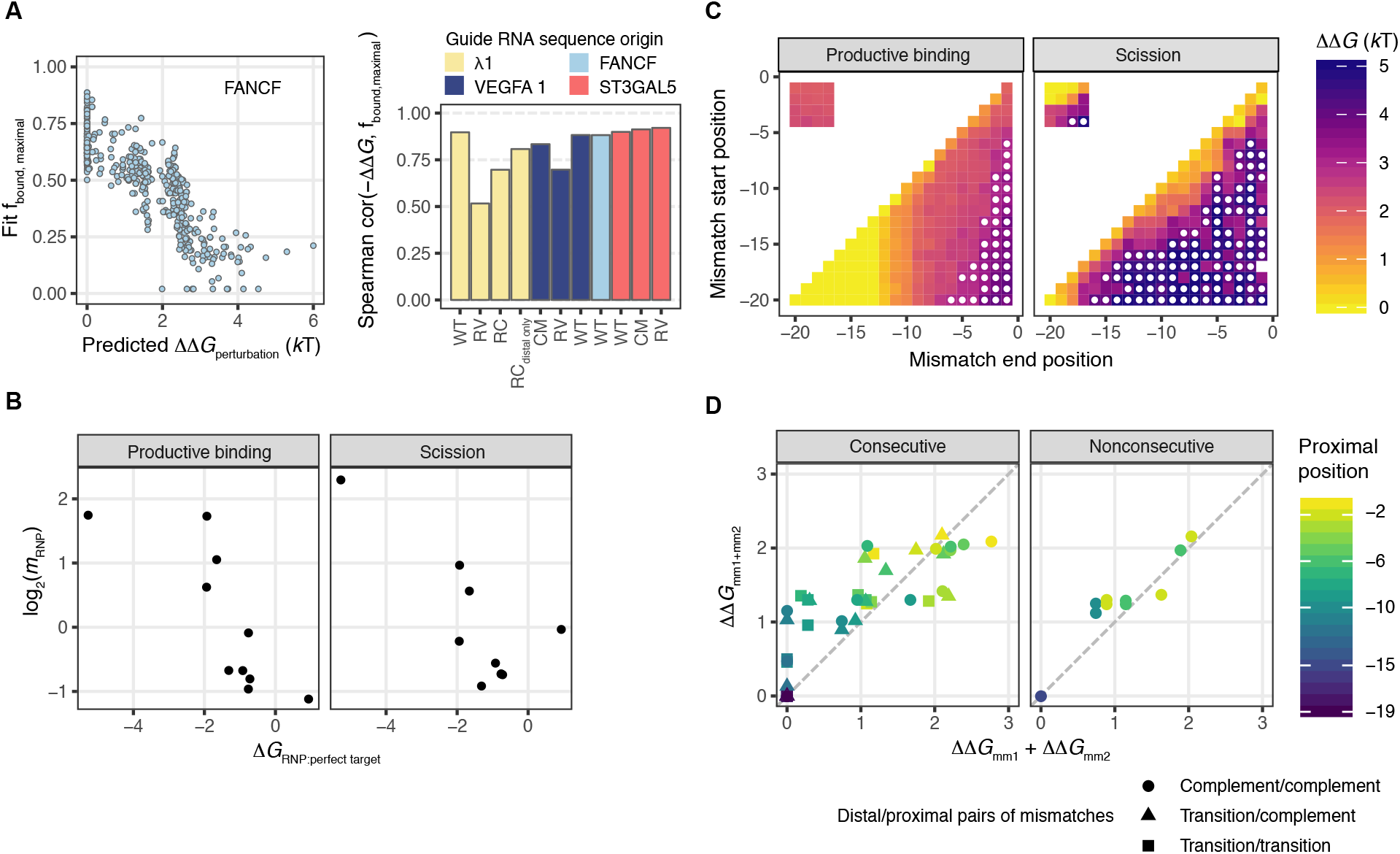
A) Results of leave-one-out cross-validation of fit biophysical parameters. Performance was assessed by Spearman correlation between fit *ΔΔG*_perturbation_ and left-out f_bound, maximal_. FANCF data is shown on left, correlations for all dCas9 RNPs on right. B) Scatterplot of *G*_RNP:perfect target_ against log2-transformed *m*_RNP_ values. C) Fit *ΔΔG*_perturbation_ values for mismatch series targets for productive binding and scission. Complement mismatches (e.g. rA:dA) start at the position on the y-axis and continue to the position on the x-axis. Tiles in the top left corner wrap from PAM-proximal positions to the most PAM-distal (e.g. mismatches at position −2, −1, −20 and −19). D) Fit *ΔΔG*_perturbation_ values for consecutive and programmed nonconsecutive double mismatches, visualized by the sum of their constitutive single mismatches against their fitted value. Consecutive mismatches vary by the DNA bases of the mismatches, reported as distal/proximal pairs. Color shows the location of the proximal mismatch.

**Figure S6:**
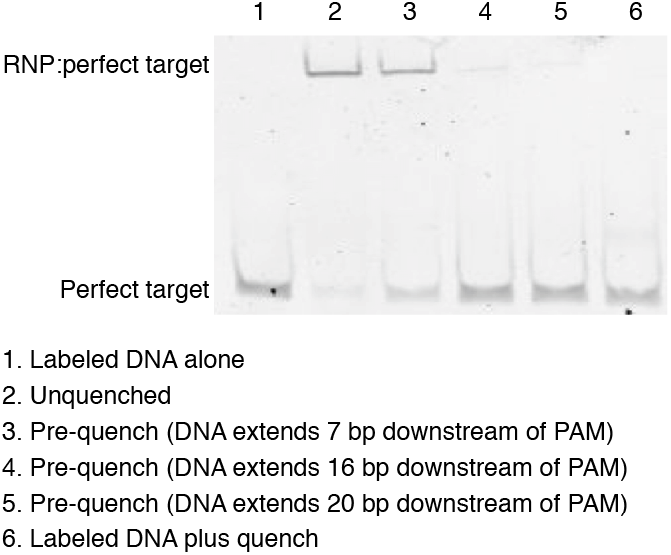
EMSA results for preemptively quenching λ1 perfect target dCas9 RNP association with different length unlabeled substrates shows the ability of different length DNA competitors to quench dCas9 binding capacity.

## References and Notes

1. H. Wang, M. La Russa, L. S. Qi, CRISPR/Cas9 in Genome Editing and Beyond. Annu. Rev. Biochem. (2016), doi:10.1146/annurev-biochem-060815-014607.

2. M. Adli, The CRISPR tool kit for genome editing and beyond. Nat Commun. 9, 1911–13 (2018).

3. J. G. Doench et al., Optimized sgRNA design to maximize activity and minimize off-target effects of CRISPR-Cas9. Nat. Biotechnol. 34, 184–191 (2016).

4. J. Listgarten et al., Prediction of off-target activities for the end-to-end design of CRISPR guide RNAs. Nat Biomed Eng. 2, 38–47 (2018).

5. D. Singh, S. H. Sternberg, J. Fei, J. A. Doudna, T. Ha, Real-time observation of DNA recognition and rejection by the RNA-guided endonuclease Cas9. Nat Commun. 7, 12778 (2016).

6. H. Nishimasu et al., Crystal structure of Cas9 in complex with guide RNA and target DNA. Cell. 156, 935–949 (2014).

7. F. Jiang et al., Structures of a CRISPR-Cas9 R-loop complex primed for DNA cleavage. Science. 351, 867–871 (2016).

8. M. Jinek et al., Structures of Cas9 endonucleases reveal RNA-mediated conformational activation. Science. 343, 1247997–1247997 (2014).

9. J. S. Chen et al., Enhanced proofreading governs CRISPR-Cas9 targeting accuracy. Nature. 550, 407–410 (2017).

10. D. Singh et al., Mechanisms of improved specificity of engineered Cas9s revealed by single-molecule FRET analysis. Nat. Struct. Mol. Biol. 25, 347–354 (2018).

11. M. D. Newton et al., DNA stretching induces Cas9 off-target activity. Nat. Struct. Mol. Biol. 26, 185–192 (2019).

12. S. Gong, H. H. Yu, K. A. Johnson, D. W. Taylor, DNA Unwinding Is the Primary Determinant of CRISPR-Cas9 Activity. Cell Rep. 22, 359–371 (2018).

13. A. T. Raper, A. A. Stephenson, Z. Suo, Functional Insights Revealed by the Kinetic Mechanism of CRISPR/Cas9. J. Am. Chem. Soc. 140, 2971–2984 (2018).

14. C. Jung et al., Massively Parallel Biophysical Analysis of CRISPR-Cas Complexes on Next Generation Sequencing Chips. Cell. 170, 35–47.e13 (2017).

15. E. A. Boyle et al., High-throughput biochemical profiling reveals sequence determinants of dCas9 off-target binding and unbinding. Proc. Natl. Acad. Sci. U.S.A. 114, 5461–5466 (2017).

16. R. Nutiu et al., Direct measurement of DNA affinity landscapes on a high-throughput sequencing instrument. Nat. Biotechnol. 29, 659–664 (2011).

17. T. Schneider et al., CircRNA-protein complexes: IMP3 protein component defines subfamily of circRNPs. Sci Rep. 6, 31313–11 (2016).

18. A. Zykovich, I. Korf, D. J. Segal, Bind-n-Seq: high-throughput analysis of in vitro protein-DNA interactions using massively parallel sequencing. Nucleic Acids Research. 37, e151–e151 (2009).

19. M. Levo et al., Unraveling determinants of transcription factor binding outside the core binding site. Genome Res. 25, 1018–1029 (2015).

20. H. Xu et al., Sequence determinants of improved CRISPR sgRNA design. Genome Res. 25, 1147–1157 (2015).

21. M. A. Horlbeck et al., Compact and highly active next-generation libraries for CRISPR-mediated gene repression and activation. Elife. 5(2016), doi:10.7554/eLife.19760.

22. X. Wu et al., Genome-wide binding of the CRISPR endonuclease Cas9 in mammalian cells. Nat. Biotechnol. 32, 670–676 (2014).

23. V. Pattanayak et al., High-throughput profiling of off-target DNA cleavage reveals RNA-programmed Cas9 nuclease specificity. Nat. Biotechnol. 31, 839–843 (2013).

24. R. Lorenz et al., ViennaRNA Package 2.0. Algorithms Mol Biol. 6, 26–14 (2011).

25. S. B. Thyme, L. Akhmetova, T. G. Montague, E. Valen, A. F. Schier, Internal guide RNA interactions interfere with Cas9-mediated cleavage. Nat Commun. 7, 11750 (2016).

26. M. Jost et al., Titrating gene expression using libraries of systematically attenuated CRISPR guide RNAs. Nat. Biotechnol. 6, e1001154–10 (2020).

27. S. H. Sternberg, B. LaFrance, M. Kaplan, J. A. Doudna, Conformational control of DNA target cleavage by CRISPR-Cas9. Nature. 527, 110–113 (2015).

28. Y. S. Dagdas, J. S. Chen, S. H. Sternberg, J. A. Doudna, A. Yildiz, A conformational checkpoint between DNA binding and cleavage by CRISPR-Cas9. Sci Adv. 3, eaao0027 (2017).

29. J. G. Doench et al., Rational design of highly active sgRNAs for CRISPR-Cas9-mediated gene inactivation. Nat. Biotechnol. 32, 1262–1267 (2014).

30. S. Oehler, R. Alex, A. Barker, Is nitrocellulose filter binding really a universal assay for protein-DNA interactions? Anal. Biochem. 268, 330–336 (1999).

31. N. Bisaria, I. Jarmoskaite, D. Herschlag, Lessons from Enzyme Kinetics Reveal Specificity Principles for RNA-Guided Nucleases in RNA Interference and CRISPR-Based Genome Editing. Cell Syst. 4, 21–29 (2017).

32. A. Vigouroux, E. Oldewurtel, L. Cui, D. Bikard, S. van Teeffelen, Tuning dCas9’s ability to block transcription enables robust, noiseless knockdown of bacterial genes. Mol. Syst. Biol. 14, e7899 (2018).

33. X. Zhu et al., Cryo-EM structures reveal coordinated domain motions that govern DNA cleavage by Cas9. Nat. Struct. Mol. Biol. 26, 679–685 (2019).

34. J. K. Lee et al., Directed evolution of CRISPR-Cas9 to increase its specificity. Nat Commun. 9, 3048–10 (2018).

35. I. M. Slaymaker et al., Rationally engineered Cas9 nucleases with improved specificity. Science. 351, 84–88 (2016).

36. S. K. Jones et al., Massively parallel kinetic profiling of natural and engineered CRISPR nucleases. bioRxiv, 696393 (2019).

37. J. H. Hu et al., Evolved Cas9 variants with broad PAM compatibility and high DNA specificity. Nature. 556, 57–63 (2018).

38. S. Q. Tsai et al., CIRCLE-seq: a highly sensitive in vitro screen for genome-wide CRISPR-Cas9 nuclease off-targets. Nat Methods. 14, 607–614 (2017).

39. P. Cameron et al., Mapping the genomic landscape of CRISPR-Cas9 cleavage. Nat Methods. 14, 600–606 (2017).

40. Q. Zhang et al., The post-PAM interaction of RNA-guided spCas9 with DNA dictates its target binding and dissociation. Sci Adv. 5, eaaw9807 (2019).

41. L. A. Gilbert et al., CRISPR-mediated modular RNA-guided regulation of transcription in eukaryotes. Cell. 154, 442–451 (2013).

42. J. Tycko et al., Mitigation of off-target toxicity in CRISPR-Cas9 screens for essential non-coding elements. Nat Commun. 10, 4063–14 (2019).

43. T. Wang et al., Pooled CRISPR interference screening enables genome-scale functional genomics study in bacteria with superior performance. Nat Commun. 9, 2475–15 (2018).

44. F. Rousset et al., Genome-wide CRISPR-dCas9 screens in E. coli identify essential genes and phage host factors. PLoS Genet. 14, e1007749 (2018).

45. J. C. Klein et al., Multiplex pairwise assembly of array-derived DNA oligonucleotides. Nucleic Acids Research. 44, e43–e43 (2016).

46. M. Sturm, C. Schroeder, P. Bauer, SeqPurge: highly-sensitive adapter trimming for paired-end NGS data. BMC Bioinformatics. 17, 208–7 (2016).

47. T. Magoč, S. L. Salzberg, FLASH: fast length adjustment of short reads to improve genome assemblies. Bioinformatics. 27, 2957–2963 (2011).

